# Conscious processing of narrative stimuli synchronizes heart rate between individuals

**DOI:** 10.1101/2020.05.26.116079

**Authors:** Pauline Pérez, Jens Madsen, Leah Banellis, Başak Türker, Federico Raimondo, Vincent Perlbarg, Melanie Valente, Marie-Cécile Niérat, Louis Puybasset, Lionel Naccache, Thomas Similowski, Damian Cruse, Lucas C Parra, Jacobo D Sitt

**Author notes:** First authors contributed equally (Pauline Pérez, Jens Madsen). Last authors contributed equally (Lucas C Parra, Jacobo D Sitt).

## Abstract

Heart rate has natural fluctuations that are typically ascribed to autonomic function. Recent evidence suggests that conscious processing can affect the timing of the heartbeat. We hypothesized that heart rate is modulated by conscious processing and therefore dependent on attentional focus. To test this, we leverage the observation that neural processes can be synchronized between subjects by presenting an identical narrative stimulus. As predicted, we find significant inter-subject correlation of the heartbeat (ISC-HR) when subjects are presented with an auditory or audiovisual narrative. Consistent with the conscious processing hypothesis, we find that ISC-HR is reduced when subjects are distracted from the narrative, and that higher heart rate synchronization predicts better recall of the narrative. Finally, patients with disorders of consciousness who are listening to a story have lower ISC-HR, as compared to healthy individuals, and that individual ISC-HR might predict a patients’ prognosis.. We conclude that heart rate fluctuations are partially driven by conscious processing, depend on attentional state, and may represent a simple metric to assess conscious state in unresponsive patients.

## Introduction

In healthy individuals, heart rate fluctuates with breathing and changes in parasympathetic and sympathetic tone ^1–3^. Physical activity naturally increases heart rate, but also just thinking about physical activity may increase heart rate^4^. Similarly, mental exercises such as meditation can reduce heart rate^5^. The effect of cognition on heart rate is perhaps even more direct than these traditional accounts ^6,7^. We also know that suspense and surprise can transiently increase heart rate^8^. Most likely these immediate effects of the mind on the heart subserved the purpose of preparing the body for imminent action^9^. Despite this evidence, the role of (un)conscious perception^10^ on heart rate is less clear. It is well established that the brain can unconsciously detect novelty in the stimulus, as demonstrated with event-related potential studies (e.g. MMN^11–13^, and N400^14,15^). Recent evidence shows that the timing of an individual heartbeat may be affected by the perception of an unexpected sound, but only when consciously perceived^16^. We hypothesize that conscious processing of perceptual information will affect heart rate. Therefore, we predict that fluctuations in heart rate will depend on top-down attention to the stimulus and predict memory performance, a known factor^17^ and a correlate of conscious perception.^18^

To test these predictions, we will leverage the observation that natural narrative stimuli guide cognitive processes resulting in reliable neural responses. This was first observed by measuring hemodynamic brain activity during movies: when humans watch the same movie, they have similar fluctuations in brain blood oxygenation^19^. Specifically, the temporal fluctuations of the signal measured with functional magnetic resonance (fMRI) are correlated between subjects. Significant inter-subject correlation of brain activity has now been observed with other neuroimaging modalities, including EEG, MEG and fNIRS^10–13^. Thus, neurophysiological fluctuations appear to synchronize on a wide range of time scales, from milliseconds to several minutes. This phenomenon is also not constrained to movies but has been observed for speech, music, or during driving ^20–22^. There are even significant correlations in the spatial patterns of fMRI activity between speakers and listeners^23^ or the time-courses of EEG signals of two individuals engaged in a conversation^24^. This similarity of neural activity in response to narrative stimuli suggests that these stimuli elicit similar perceptual and cognitive processes in different subjects.

Consistent with this, inter-subject correlation crucially depends on the cognitive state of the participant. Subjects that are not attentive or do not follow the narrative show significantly reduced inter-subject correlation, both in EEG and fMRI ^25–27^. A drop in inter-subject correlation is also observed in patients with disorders of consciousness relative to healthy controls ^28–30^. Indeed, a cohesive narrative is crucially important to elicit synchronized brain activity in fMRI, in particular at long time scales.^31^ It comes as no surprise then that inter-subject correlation has been found to be predictive of a variety of behavioral outcomes, such as audience retention, memory of content, efficacy of advertising, efficacy of communication and political speeches, and more ^23,32–36^.

There are many studies reporting a correlation of physiological signals across subjects.^37^ Generally this has been linked to physical or social interaction,^38–40^ or at the very least a copresence at the same place and time.^41^ However, consistent with our hypothesis, the simultaneous experience is not crucial for synchronization. A few recent studies report a correlation of heart rate fluctuations across subjects watching the same movie at different times, and ascribe this to shared emotions elicited by the film.^42,43^

Our conscious processing hypothesis predicts that this synchronization phenomenon will occur not just for the film, but more generally for narrative stimuli, that inter-subject correlation of heart rate will be modulated by attention, that it will correlate with cognitive performance, and more dramatically, that it will be reduced in patients with disorders of consciousness. We confirm these predictions in a series of four experiments and conclude that heart rate synchronization has the potential to become a marker of cognitive state in a clinical setting.

## Results

In all four experiments we presented narrative stimuli to each subject while recording their electrocardiogram (EKG, Figure 1A), in experiments 3 and 4 we also recorded respiratory activity. Recordings were aligned in time between subjects and instantaneous HR was estimated as the inverse of the RR intervals (Figure 1B). Mean and standard deviation of these instantaneous measures provide HR and HR variability (HRV) for each subject. The instantaneous HR signals are upsampled to a common sampling rate and correlated between all pairs of subjects (Figure 1C). Inter-subject correlation of HR (ISC-HR) is then defined for each subject as the average Pearson’s correlation with all other subjects (Figure 1D).

**Figure 1:**
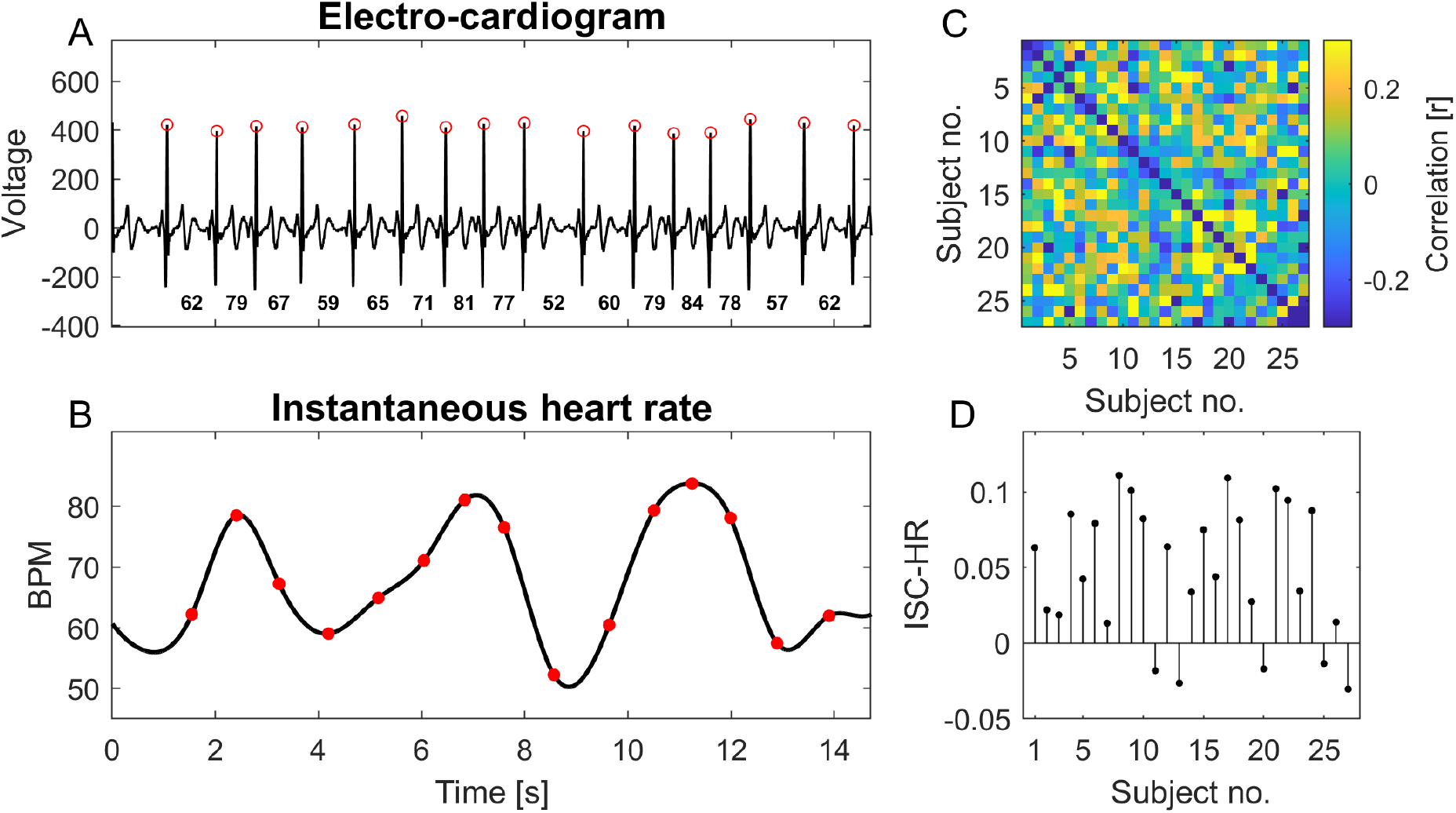
Inter subject correlation of heart rate (ISC-HR) **A:** Electro-cardiogram with peak of the R-wave detected (red o). **B:** The inverse of the interval between two R-waves defines the instantaneous heart rate (red o). This is interpolated (black) to convert heart rate into a signal with a uniform sampling rate across subjects. **C:** Pearson’s correlation coefficient of this instantaneous heart rate between pairs of subjects. **D:** Inter-subject correlation of heart rate (ISC-HR) is computed for each individual as the mean across a row of this correlation matrix. Example in this figure is taken from history segment #1, dataset #1.

### Auditory narratives synchronize listeners’ heart rate fluctuations

The objective of the first experiment was to determine whether a common auditory narrative elicits similar heart rate fluctuations in healthy volunteers (Experiment 1). Subjects were presented with one-minute segments of an audiobook of Jules Verne’s “20,000 leagues under the sea”. First we tested whether there was significant inter-subject correlation of the instantaneous HR. To this end we compared the ISC-HR values to values computed on signals randomly shifted in time within-subjects (see methods). When this analysis is performed on individual one-minute segments, only a few subjects show significant non-zero ISC-HR (Fig. 2A, black dots, FDR at 0.05). When averaging ISC-HR values over the 16 minutes, 17 of the 27 subjects show statistically significant HR correlation (Fig. 2B; FDR at 0.05). No significant negative correlations were found. As an additional control, we randomly shuffle the one-minute story segments between subjects breaking the narrative synchrony across subjects. As expected, we observed a significant drop in ISC values between the original and permuted conditions (Fig. 2C, paired t-test (26) = 9.20, p = 2. 10-9). In total, we conclude that the narrative stimulus induces similar HR fluctuations across subjects. ISC-HR therefore captures how strongly the stimulus drives the fluctuations of HR in each subject.

Results on average HR and HRV and their potential relation to ISC-HR are generally unremarkable for these data and are discussed in the Supplement (Fig. S1).

**Figure 2:**
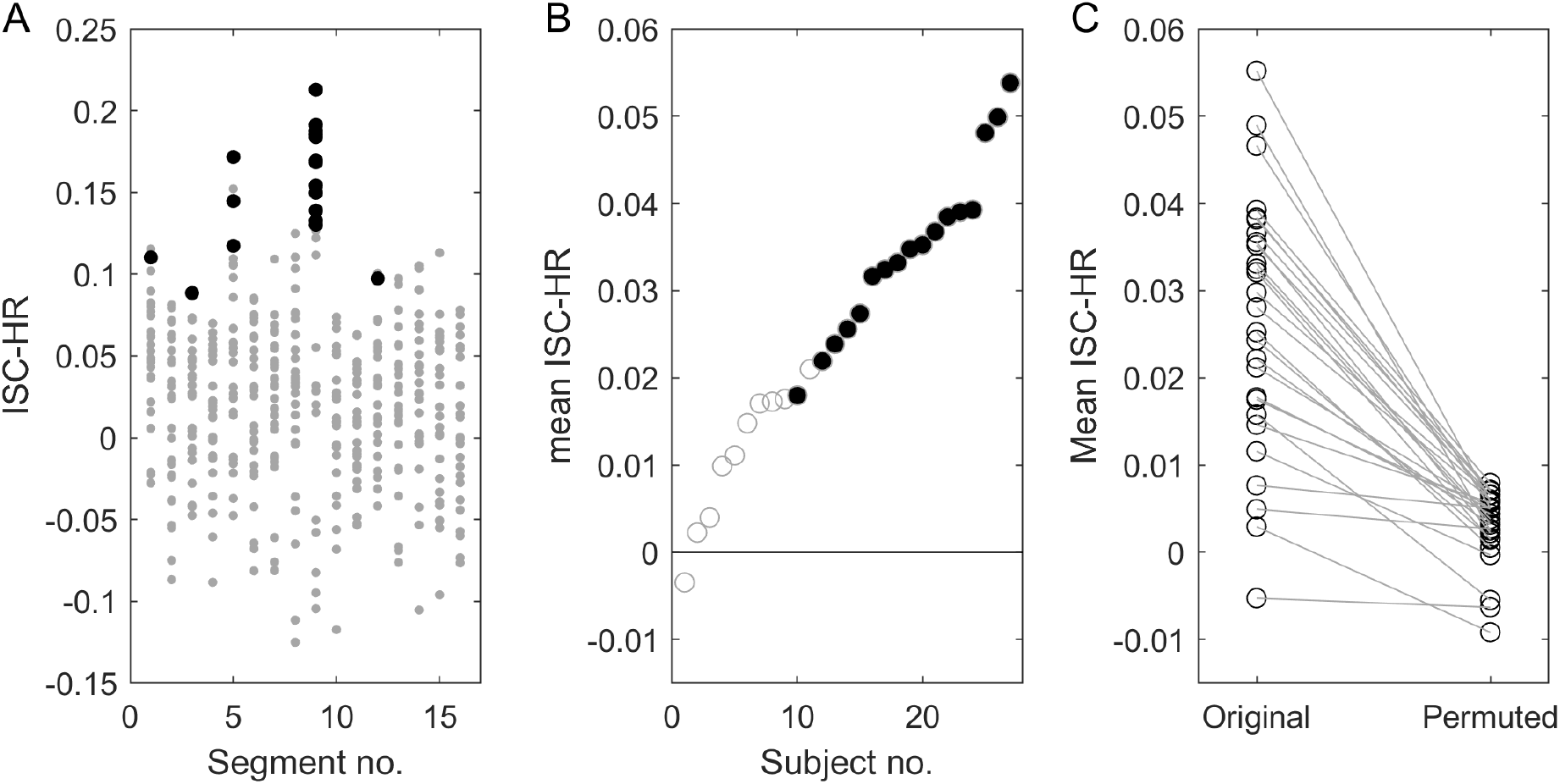
ISC-HR resolved in time and by subject. In Experiment 1, subjects listened to segments of audio narratives of 60 seconds each (N=16). **A**: ISC-HR is computed for each subject (N=27) and each segment. **B:** For each of the 27 subjects ISC is averaged over the 16 segments. Subjects are ordered by their ISC values. Black points (panel A and B) indicate statistically significant ISC values. Gray points are not statistically significant. Statistical significance is determined using circular shuffle statistics (10,000 shuffles and corrected for multiple comparisons with FDR of 0.01). Specifically, the heart rate signal of each subject is randomly shifted in time. **C:** As additional control here ISC is compared to the ISC obtained when story segments are swapped across subjects at random.

### Attention modulates synchronization of HR fluctuations during audiovisual narratives

We demonstrated above that an auditory narrative can synchronize HR fluctuations across subjects. In the second experiment we aim to determine if this synchronization is modulated by attention to the stimulus (Experiment 2). Here we used short and engaging instructional videos of 3-5 minute duration, similar to our previous work^26^. Each subject viewed 5 videos in sequence normally. Then they view the same videos a second time, but now with the instruction to count backward silently in their mind in step 7 starting from a random number. This secondary task aims to distract subjects from viewing the video ^25,26^.

We find that ISC-HR drops in the distracted condition relative to the normal attentive state (Fig. 3A). A repeated-measures ANOVA shows a strong effect of attention (F(1,238)=73.45, p=1.32e-15) and an effect of the video (F(4,238)=14.59, p=1.14e-10) as well as a subject effect (F(26,238)=2.86, p=1.34e-05). The effect of attention is significant for each story individually (follow up pairwise t-test, all p<0.05) and when averaging over all 5 videos with a total duration of 22:33 minutes we see a numerical drop in ISC-HR with distraction in all but one of the 27 subjects (Fig. 3B).

**Figure 3:**
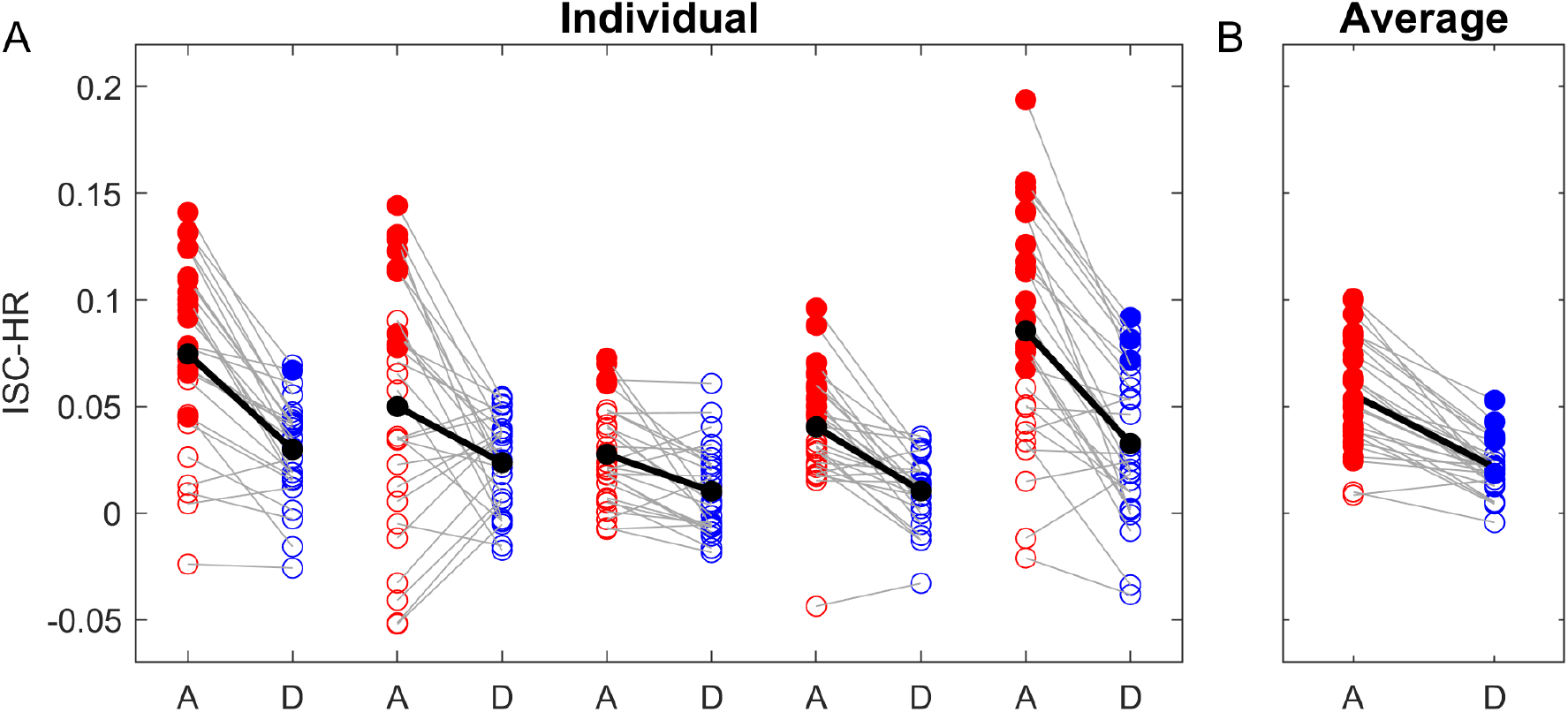
Intersubject correlation (ISC) of the instantaneous heart rate is modulated by attention. In Experiment 2, 27 subjects watched 5 educational videos of 3-5 minute duration each. Here ISC is measured against the attentive condition, i.e. both attentive and distracted subjects are correlated against the HR collected during the attentive condition. Filled points indicate individually statistically significant ISC-HR (FDR <0.01). **A:** Subjects watched the same videos twice, either in an attentive (A, red) or distracted (D, blue) condition. ISC was systematically higher in the attentive condition for the five videos. Gray lines indicate individual subjects and the black lines the group average. **B:** Same results when average across the five videos.

### Attention modulates HRV, but this is not the driving factor in modulation of ISC

For this Experiment 2, in addition to ISC we also analyzed heart rate variability (HRV), defined here as the standard deviation of instantaneous HR (Fig. S2B). We see an increase in HRV when subjects are distracted. An ANOVA shows a fixed effect of attention (F(1,211)=63.60, p=9.54e-14), random effect of subject (F(23,211)=29.56, p=1.57e-53), but we see no significant video effect (F(4,211)=0.77, p=5.44e-01). Perhaps the increase in HRV in the distracted condition could explain the drop in ISC-HR. If this was the case, we would expect that HRV correlates negatively with ISC-HR as, by definition, the two are inversely related. However, the opposite seems to be the case: subjects with higher HRV have also higher ISC-HR (Fig. S2D). Therefore, it appears that the modulation of HRV and ISC-HR are independent phenomena. The effects on mean HR were generally unremarkable (Fig. S2A & S2C).

### Synchronization of HR fluctuations is modulated in the time scale of 5-10 seconds

It is well established that during waking rest HR fluctuates at various timescales ^44^. This is reproduced in the present context of video presentation (of Experiment 2) by computing HRV after band-pass filtering the instantaneous HR in different frequency bands (Fig. 4A). To determine which time scale dominates ISC and its modulation with attention we computed ISC similarly resolved by frequency band (Fig. 4B). We find that ISC and its modulation with attention are dominant in the low-frequency range from 0.09 Hz to 0.15 Hz (p < 0.00001 as a single cluster). It is worth noting that ISC was not modulated by attention in the high-frequency peak (around 0.3Hz), which corresponds to the dominant frequency of breathing (Supplementary Fig. S5).

**Figure 4:**
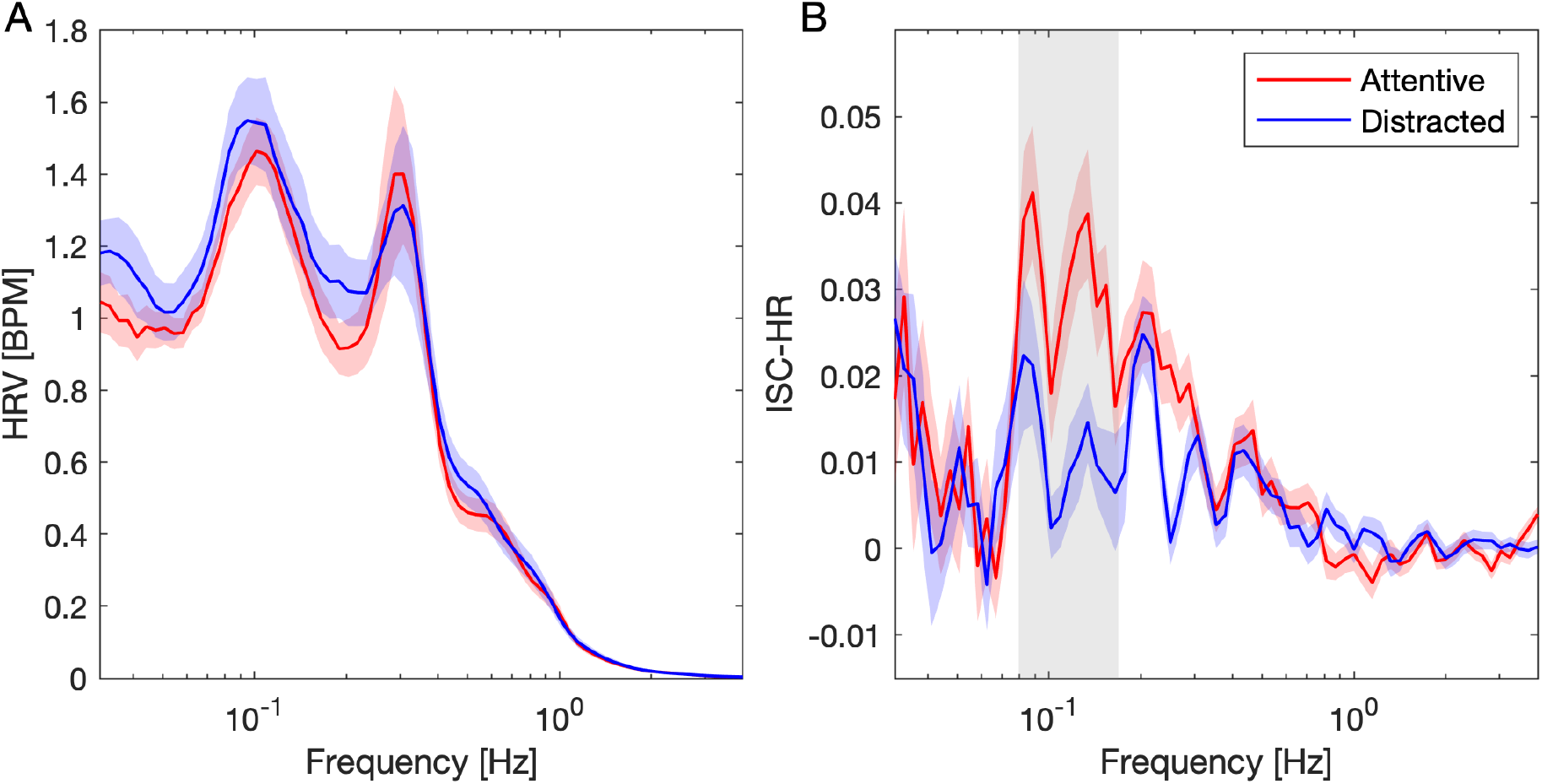
Spectrum of instantaneous HR and ISC-HR and its modulation with attention. For Experiment 2, instantaneous HR was band-pass filtered with center frequency on a logarithmic scale and a bandwidth of 0.2 of the center frequency. **A:** HRV is computed here as the root mean square of the band-passed instantaneous HR averaged over the 5 videos (~15 min total). The greyed area around the curves in both panels is the standard error. **B:** ISC-HR is computed as before, but now on the band-passed instantaneous HR and averaged over the 5 videos. In both panels significant differences between attending and distracted conditions are established in each band with a paired t-test over the 27 subjects (multiple comparison corrected with one-dimensional cluster statistics, p<0.05). Colored-shaded areas indicate standard error of the mean.

### Attention modulates synchronization of HR fluctuations during audio-only narratives, but does not synchronize breathing

Given the dependence of HR fluctuations on attention, we expected that HR would be predictive of cognitive processing of the narrative. In the next experiment (Experiment 3) we, therefore, recorded HR during the presentation of auditory narratives, and afterward asked adult subjects to recall factual information presented in the story, e.g. “What was the name of the two main characters?”. Subjects listened to four auditory narratives, in either an attentive or distracted condition. This time the narratives were children’s stories of 8-11 minute duration, and the secondary task consisted of counting target tones that were inserted in the audio asynchronously across subjects. To rule out order effects we now divided the participants in two groups. In group 1 (N=9) subjects listened to stories 1 and 2 in the attentive condition and stories 3 and 4 in the distracted condition. In group 2 (n=12) subject listened to the same stories with the attention condition reversed.

We find again that ISC-HR drops significantly when subjects are distracted (Fig. 5A, paired t-test t(20)=8.2, p< 10^-7^). Similarly to experiment 2 with video, here 15 out of the 21 subjects show a statistically significant correlation of HR in the attentive condition and none in the distracted condition. As expected, subjects performed significantly better in recalling elements of the story in the attentive condition as compared to the distracted condition (Supplementary Fig. S3, Wilcoxon Signed-Rank Test, z = 4.03, p = 5.7e-5).Our hypothesis postulates that ISC-HR is the result of similar conscious processing of the narrative stimulus, thus, we predicted that subjects with higher ISC-HR will be better at remembering elements of the stories. Indeed, we find that ISC correlates with memory recall performance across conditions (Fig 5B, r(40)=0.767, p=3.1e-9, Spearman’s correlation is used here due to the bounded nature of the percent measure). More importantly, even within the normally attentive condition with a normal fluctuation of HR we find that ISC-HR is predictive of memory performance (r(19)=0.57, p=7.3 e-3). In the distracted conditions there was no correlation with memory performance (r(19)=-0.1, p=0.67, BF01 = 4.88) possibly because ISC-HR was not statistically significant for any of the subjects. Overall, we conclude that ISC-HR is indicative of conscious processing of the narrative.

**Figure 5:**
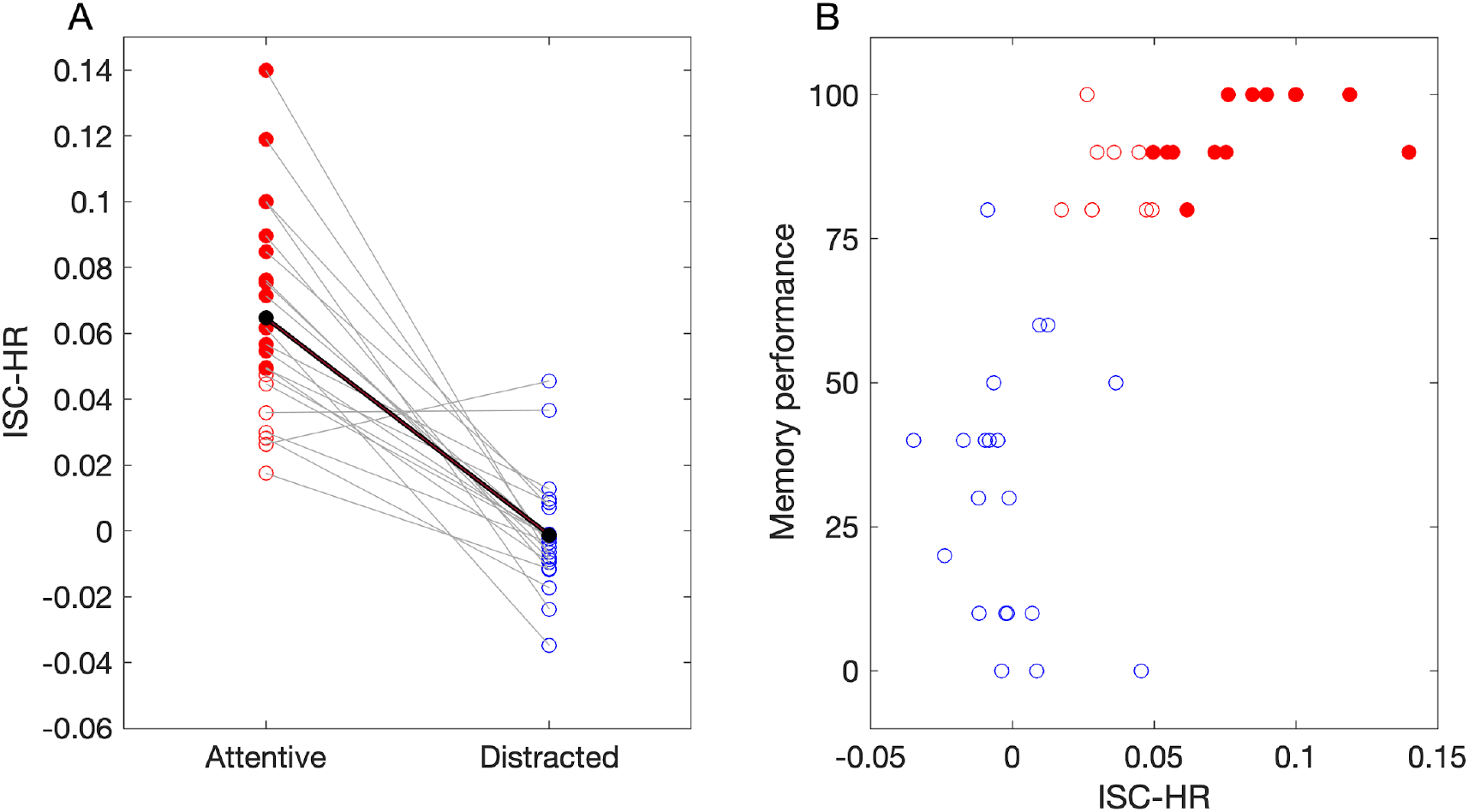
Intersubject correlation is higher when subjects were attentive to the auditory narrative and this correlation indexes the subjects’ memory performance. In Experiment 3, Subjects listened to four recordings of children’s stories 8-10 minutes in duration. Subjects were instructed to either attend to a story normally (attentive, red), or to count backward when they heard a target sound inserted in the audio (distracted, blue). Again, ISC is measured against the attentive condition, i.e. both attentive and distracted subjects are correlated against the HR collected during the attentive condition. Filled points indicate individually statistically significant ISC-HR (FDR <0.01). **A:** ISC-HR for each subject (N=21) averaged over four stories. **B:** Memory performance measured as percent of correct answers to free recall questions about the content of the stories. Filled and empty circles indicate significant and nonsignificant ISC-HR respectively (p<0.05 shuffle statistics)

### Intersubject correlation of heart rate is not driven by synchronous breathing

It is well established that HR fluctuations are driven in part by breathing.^45^ This phenomenon is known as respiratory sinus arrhythmia and can affect a range of frequencies ^46,47^. It is possible that the attentional modulation in this frequency band is caused by synchronization of breathing between subjects. We, therefore, collected in Experiments 3 respiratory movement concurrently with the EKG and measured inter-subject correlation of breathing. First, we tested if the power spectrum of the raw respiratory signal changes with attention and found small increases in power at “high” frequencies (above 0.3Hz, Supplementary Fig. S5). Second, we validated the relationship between respiratory and cardiac activity by computing the correlation between respiratory signal and instantaneous HR. In the attentive condition twelve (out of 21) subjects showed a significant cardio-respiratory coupling, and in the distracted condition it was fifteen (out of 21) (Supplementary Fig. S6A). In addition, we found a non-significant reduction in breathing-HR correlation in the distracted condition versus the attentive (Fig S6A, t(20)=2.0, p=0.18 (FDR corrected), BF10 = 1.2, BF01 = 0.83). Third, we tested for significant ISC in the raw breathing signal and various of its features, specifically, the instantaneous respiratory frequency and amplitude computed separately for inspiration and expiration (using the breathmetrics toolbox ^48^). We did not find a significant ISC of the raw breathing signal in any of the subjects, nor was there any effect of attention when comparing across all subjects (Fig. S6B, paired t-test, t(20)=-0.8, p=0.75 (FDR corrected), BF01 = 3.21). Similarly, none of the respiration features showed significant ISC, nor a drop in ISC for the distracted conditions (Supplementary Fig S6C-F). In other words, the auditory narratives did not reliably entrain the subjects breathing nor was this modulated by attention. On the flipside, while Bayes factor analysis provides some evidence for a lack of synchronized breathing, the evidence is only moderate (in all instances 1<BF<4 in favor of this Null hypothesis, Supplementary Fig S6).

Finally, to determine if delayed influences of respiration could explain the results obtained from HR, for each subject and condition we subtracted from the HR signals any instantaneous or time delayed linear correlation of the respiratory signal and recomputed the ISC-HR. We obtained similar modulation of ISC with attention (Supplementary Fig S7, t-test: t(20) = 5.57, p = 1.9e-05, BF = 1251.3).

In conclusion, while we do not have strong evidence against a synchronization of breathing, these results do suggest that the effect of cognition on synchronizing HR cannot be explained in this study by the synchronization of breathing.

### Synchronization of heart rate fluctuations is disrupted in patients with disorder of consciousness

Given the dependence on attention and conscious processing of these synchronized HR fluctuations, we predicted that patients with disorders of consciousness (DOC) will have diminished HR synchronization when presented with an auditory narrative. We recorded EKG in 19 DOC patients, in addition to 24 healthy controls (Experiment 4). The patients were hospitalized to determine their state of consciousness and neurological prognosis. Patients were behaviorally assessed using the standard Coma Recovery Scale-revised^49^. State of consciousness was determined using the currently accepted categorization^50^, patients were classified either in (1) Coma, (2) Vegetative state/Unresponsive wakefulness syndrome (UWS), (3) Minimally conscious state minus (MCS-), (4) Minimally conscious state plus (MCS+), and exit Minimally conscious state (EMCS) (See Supplementary Table 3 a detailed description of the patients). Patients and healthy subjects listened through headphones to a children’s story of 10-minute duration. Using the data from healthy subjects we first replicated the results of Experiment 3 showing that ISC computed for HR is systematically larger than the ISC of any of the respiratory features tested (Supplementary Fig. S8). We then computed ISC-HR by correlating HR with that of healthy controls. As expected ISC-HR values were lower in patients (Fig 6A, t-test: t(41) = 3.14, p = 0.003, BF10 = 12.4). Within patients no significant correlation was found between ISC-HR and state-of-consciousness (Fig. 6A, Spearman correlation, R(17) = −0.28, p = 0.24, BF01 = 4.07) nor between ISC-HR and the Coma Recovery Scale-Revised (CRS-R^49^; Fig 6B, Spearman correlation, R(17) = −0.3, p = 0.22, BF01 = 3.36). Reduced HRV is sometimes found in traumatic brain injury patients.^51^ We therefore analyzed HRV to verify that the drop in ISC-HR is not a noise-floor effect, i.e. if HRV drops in patients it may be difficult to measure inter-subject correlation above random fluctuations (Fig. S9B). Contrary to what was expected we found higher HRV in the DOC patients compared to healthy controls (t(41) = 2.34, p = 0.02, BF10 = 2.53), ruling out a noise-floor effect. We also found higher mean HR in DOC patients compared to healthy controls (Fig S9A; t(41) = 4.7, p = 2.9e-05, BF = 639). However, given the previous lack of correlation between ISC and mean HR we do not believe this contributed to the decrease of ISC-HR in patients.

**Figure 6:**
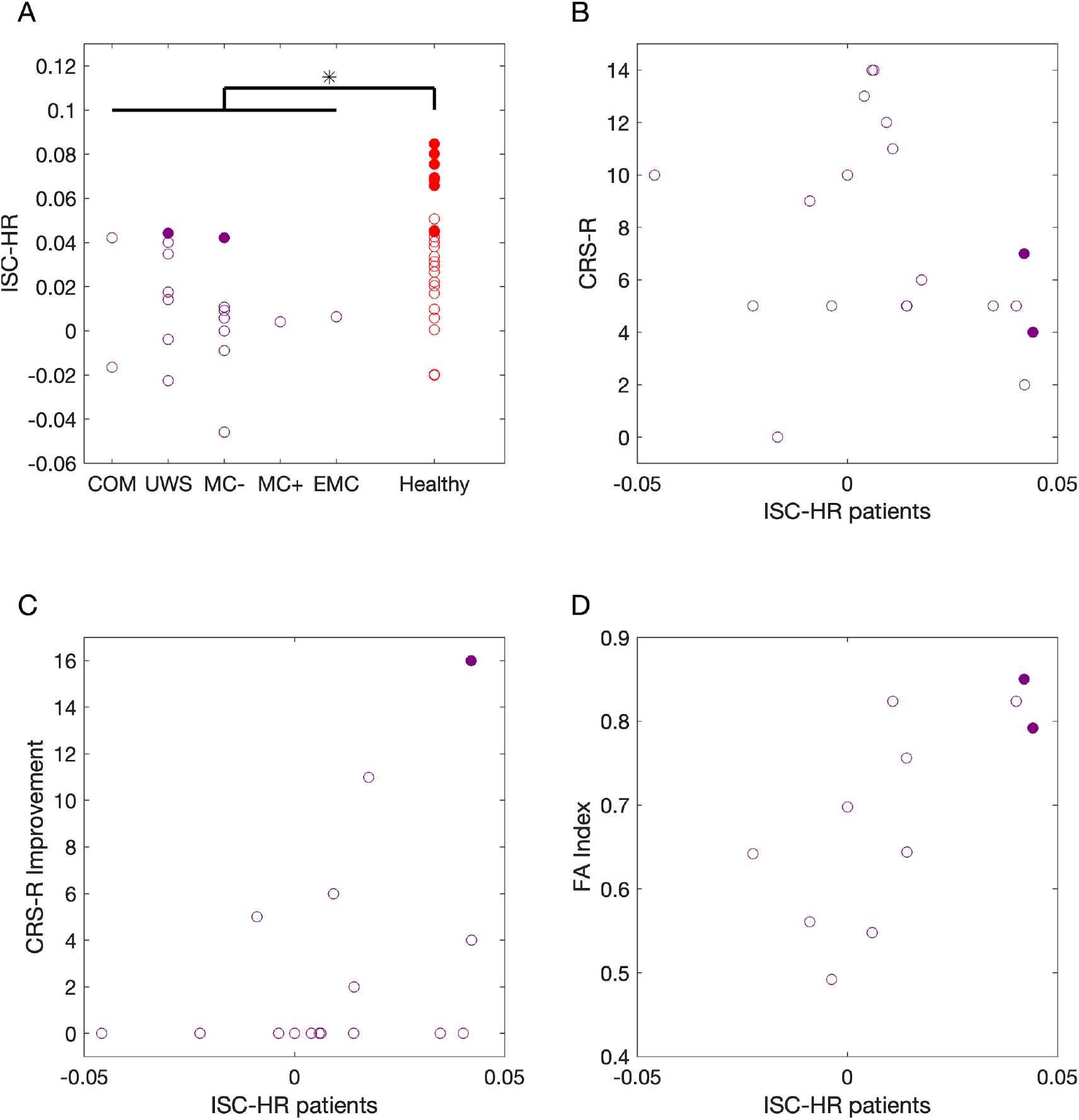
Audio narratives synchronize HR fluctuations in healthy controls but not in patients with disorder of consciousness. In Experiment 4, subjects listened to a children’s story (La part des ancêtres from Leonora Miano; 10 minutes). **A:** ISC-HR is measured by correlating instantaneous HR with that of healthy subjects. Filled cycles indicated statistically significant ISC. **B:** Comparison of the ISC-HR with Coma Recovery Scale-Revised in patients (N=19). **C:** Comparison of the ISC-HR with improvement of Coma Recovery Scale-Revised six months after the first assessment (N=17) **D:** Comparison of ISC-HR and whole-brain white matter fractional anisotropy in patients, were available (N=11).

When measured individually, only two of 19 patients showed statistically significant ISC-HR (FDR corrected p<0.05, Fig. 6, purple filled circles). For these two patients’ outcomes at the six-month follow-up were mixed; one patient fully regained consciousness whereas for the other, life-sustaining therapies were withdrawn before the follow-up assessment. Among the remaining 17 patients only one additional patient recovered consciousness, although in a completely aphasic condition. These results suggest that the patients’ ISC-HR might carry prognostic information with a specific emphasis on conscious verbal processing. To test this hypothesis, we first correlated the patients’ ISC-HR to the CRS-R improvement after six months of the initial assessment. We found a positive correlation between ISC-HR and CRS-R improvement, although not statistically significant (Fig 6C, Spearman correlation, R(14)=0.43,p=0.097, BF10 = 0.58, second assessment was available only for 16 patients). A limitation of behavioral assessment of patients is that it cannot detect covert awareness ^52,53^, a condition that can occur in up to 15% of the UWS patients^54^. Therefore, we also correlated the patients’ ISC-HR to an anatomical measure of brain integrity -- whole-brain white matter fractional anisotropy. This FA index has been linked to neurological recovery in DOC patients^55^. We found a significant correlation between ISC-HR and FA Index (Fig. 6D, Spearman correlation, R(9)=0.73,p=0.01, BF10 = 5.65, FA Index was available only for 11 patients).

## Discussion

The hypothesis that motivated this set of experiments was that conscious processing of information modulates instantaneous heart rate. This fluctuating heart rate will synchronize across subjects when presented with narrative stimuli that are processed similarly. We tested the predictions resulting from this hypothesis in a series of four experiments. In the first experiment with healthy volunteers we confirmed that heart rate fluctuations correlate between subjects for auditory narratives. In the second and third experiment we confirm that distracting the participants with a secondary task reduced this correlation, for video and audio narratives alike. Importantly, we confirm the prediction that synchronization of HR fluctuations is predictive of memory performance. We also determined that HR synchronization is likely not driven by synchronous breathing across subjects for the present audio narratives or educational videos. Finally, in the fourth experiment we presented an auditory narrative to patients with disorders of consciousness and found that their heart rate fluctuations do not correlate with that of healthy subjects. In total, we found that natural stimuli induce small but highly reliable correlations of HR, which are detectable in individual subjects and readily reproduced across four different experiments. We found a robust link between this HR synchronization and conscious processing of the audiovisual stimuli. To establish the causal direction of this link future work will require simultaneous neural recordings and prospective interventions.

There is an extensive literature demonstrating that physiological signals such as heart rate, respiration, and skin conductivity can synchronize between individuals.^37^ This literature emphasizes physical interaction and social relationships as the factors driving this synchronization. Even in the context of music, theater, or film, the emphasis is on the concurrent and shared experience of an audience that synchronizes heart rate to one-another.^38,39,56,57^ Here we have emphasized instead that it is the stimulus that synchronizes HR, or more precisely, a similar processing of a common stimulus. There is no need for individuals to directly interact, be related to one another or perceive the stimulus together at the same time. Consistent with our hypothesis, previous reports already show that emotional movies can synchronize the HR of viewers, even when watching the movie individually.^42,43^ The present findings go beyond this previous literature in that this HR synchronization phenomenon is not specific to live experiences or emotional movies. Rather, it occurs with many narrative stimuli, as we demonstrated here with audio recordings of children’s stories or animated educational videos.

There is also an extensive literature on the inter-subject correlation of brain signals evoked by dynamic natural stimuli, starting with experiments in fMRI while subjects watched movies.^19^ This work demonstrated that subjects process natural stimuli similarly, and that similarity of brain activity is predictive of memory performance.^58^ Subsequent experiments replicated these findings with EEG.^59,60^ Additionally, ISC of EEG is reduced when subjects are distracted^25^ and is reduced in patients with disorder of consciousness,^61^ similar to what we find here with the instantaneous heart rate. Given these parallels we expect that HR fluctuations will also synchronize across subjects listening to engaging music,^21^ and that HR synchronization will be a good indication of how engaging a narrative is.^33,62,63^

We suggest that some previous work on physiological synchronization of autonomic signals can be reinterpreted in the context of the present conscious processing hypothesis. For example, the same performance is judged differently depending on the social relationship of performer and audience member, suggesting that it is a different way of processing information in the audience member.^38^ In our view, it is the processing of the common stimulus, and not the co-presence in the same physical space that causes the synchronization of the heart rate fluctuations. We predict that many results obtained with live performances ^38,39,57^ or in-person interactions^26^ could be recovered with asynchronous playback of the same experience recorded with video. Evidently the experience may be less powerful than live in-person experiences,^41^ but the modulating factors of relationships, emotions, or empathy may still prevail in this virtual context.

We postulate that factors intrinsic to the story, such as semantics and emotions, drive a synchronized heart rate. This may include semantics of single-word to syntactic and multi-sentence level of representation as well as prosody, valence of single words, and more complex semantically mediated emotions. Capturing semantics and emotions require attention to the stimulus and some level of language comprehension. In this view, it is the narrative content that drives attention, engagement, interest and emotions. It is possible, indeed likely that the variations in ISC are due to this differing narrative content. Indeed we find a strong difference in ISC between stimuli, even within the same type of animated educational videos. Dependence of ISC on the stimulus has been found in previous EEG studies ^33,36,64^ and for heart rate in studies involving live performance for different pieces of classical music. ^57^ In contrast to EEG, we may expect that ISC-HR is less sensitive to low-level features of a stimulus. Neural evoked responses can be driven by low-level features such as luminance or sound fluctuations, which can elicit strong responses that would be trivially synchronized across subjects.^19,65^ To us it is less clear how such low-level stimulus fluctuations could drive HR fluctuations.

We have shown here that the effect on HR synchronization is dominant in the low frequency band around 0.1Hz, which falls in the frequency range of respiratory sinus arrhythmia.^45^ However, we show here that breathing does not synchronize between subjects during passive listening or watching, nor was respiratory sinus arrhythmia dependent on attention. While there are a variety of studies demonstrating a synchronization of breathing, most are contingent on synchronization of movement (e.g. speaking, singing, dancing)^66–68^. Studies with individual playback of video or speech have shown only very limited synchronization effects^69,70^ and to our knowledge there is no study that demonstrates that this synchronization depends on attention, consistent with our present result. Furthermore, it is worth noting that ISC was modulated by attention in the low-frequency peak of HRV but not the high-frequency peak around 0.3Hz which is the predominant breathing frequency. This parallels the observation in sleep, whereby the LF but not the HF peak is attenuated during slow-wave sleep ^44^, a period of deep sleep characterized by sensory decoupling ^71^ and a breakdown of cortical connectivity ^72^. Given the link between respiratory sinus arrhythmia and parasympathetic cardiac control,^73^ we therefore conclude that attention does not affect parasympathetic activity. One caveat to this conclusion is that respiration fluctuates on a slower time scale. It is possible that with longer recording or narrative stimuli different from those used in this study one may find a synchronization of breathing. Indeed, a framework of embodied cognition would predict a range of brain-body interactions to underlie consciousness and cognition.

To measure synchrony, we used here linear instantaneous correlation. Physiological synchrony is sometimes measured with more complex analysis methods in order to capture time-delayed or non-linear relationships ^38,74^. Such relationships may be expected in asymmetric scenarios such as an audience synchronizing with performers^38^ or mother and child ^37^. However, in the present study we have a group of participants who experience the stimulus in an identical setting and we found reliable synchronization of HR without the need to employ more complex analysis methods. For respiration, it is possible that individual subjects may have systematically delayed or differing responses compared to others. While we accounted for delayed influence of respiration on HR, we avoided more complex analysis of interaction between individual pairs of subjects, as the duration of the recordings may not support the increase in number of modeling parameters.

Finally, we made a proof-of-concept that the ISC-HR could be used as a simple marker for cognitive state in unresponsive patients. Note that this is in line with previous work on ISC of EEG^28^, fMRI^30^, and similar findings with galvanic skin response^75^. While we were able to distinguish patients from healthy controls, we were not able to resolve conscious states among patients. When contrasted to other methods (e.g. using EEG^76^, PET^77^, or fMRI^78^) this limitation - for the moment - curbs the potential use of this method to detect consciousness in patients. However, the results do suggest that the ISC-HR might carry information related to the patients’ recovery. Further validation with a larger sample of patients is needed to assess the clinical impact of the proposed method and to determine the optimal combination with other paraclinical tests. One of the limitations of this method is that it requires that the patients be conscious, but also to be able to process language. This double requirement may explain the positive test in one patient who recovered with preserved language processing, and a negative test in a patient who recovered in an aphasic condition. In addition, we believe that the 10 minutes story we used may have been too short for a reliable measure of ISC-HR. In the present experiments we required at least 15 minutes of instantaneous heart rate to detect significant ISC, and a link to memory. We suggest that future studies should use one or several narratives totaling at least 30 minutes of concurrent heart rate recordings. In addition, our results also indicate that the actual content of the story, and how engaging it is for the subject, plays a role in the individual ISC. Given the limited cognitive status of the patients it is critical to maximize this factor. We also suggest that the narratives should be adapted for every single patient, for instance by changing the name of the leading character using the patients’ own name. By doing so we will amplify the patients’ attention while keeping the overall structure of the story comparable across subjects.

## Methods

### Datasets

#### Experiment 1: auditory narratives with healthy participants

Twenty-seven native English speakers and healthy participants (22 females, age range 18-26, median 21 years old) listened to a 16-minute extract of an audiobook read by a male British English voice (20,000 leagues under the sea. Author: Jules Verne. Read by: David Linski. Public Domain (P) 2017 Blackstone Audio, Inc.) while their EKG was recorded. The audiobook extract was taken from the first chapter and half of the second chapter. The text is relatively suspenseful as it describes reports of an unknown monster that destroys ships. We divided the story into essays of approximately 1 minute each so that participants could take breaks between segments if they wished.

The instructions given to the subject were ‘to listen to the story and look at a fixation cross’. The stimuli were delivered by headphones - ER·1 Insert Earphones (Etymotic Research), using Psychopy v3.1.2. The EKG was recorded with two electrodes on the chest using SenseBox of ANT Neuro, sampled at 500Hz.

EKG data was cut into the 16 epochs of approximately 1-minute corresponding to each audio-segment.

This study was approved by the STEM ethics committee of the University of Birmingham, England.

#### Experiment 2: instructional videos with healthy participants

Thirty-one students watched 5 instructional videos while their EKG were recorded (19 females, age range 18-46, median 28 years old) in an attentive condition (A), where they were instructed to simply watch videos as they would regularly watch a video. Each educational video was 3-5 minutes long, chosen from popular YouTube channels covering biology, physics, and computer science. These are new recordings on videos we had tested previously.^36^ After the students had watched all 5 videos, they were asked to answer 10-12 questions about factual information about material conveyed in each video. Lastly, students were instructed to watch the video again in a distracted condition (D). In this condition participants were asked to silently count in their mind backwards from a random prime number above 800 and below 1000, in steps of 7.

The experiment was carried out at the City College of New York in a sound-attenuated booth. Subject wore headphones and watched the videos on a 19” monitor. The EKG was recorded with a BioSemi Active Two system at a sampling frequency of 2048Hz. 2 EKG electrodes were placed below the left collar bone and one on the left lumbar region. For segmentation of the EKG signal onset and offset triggers were used, in addition a flash and beep sound was embedded right before and after each video which were recorded using a StimTracker (Cedrus) to ensure precise alignment across all subjects. Of the thirty-one participants 4 were removed from analysis due to bad signal quality resulting in usable data for N=27 participants. The experimental protocol was approved by the Institutional Review Boards of the City University of New York. Documented informed consent was obtained from all subjects for laboratory experiments.

#### Experiment 3: auditory narratives with healthy participants and respiration

The EKG of 25 french native healthy participants (15 females; age range 22-28, median age 25 years old) listened to four stories while their EKG and respiration was recorded using a Polygraph Input Box (PIB of EGI-Geodesic’s physiological measurement system). This includes a chest belt to measure respiratory movement (Respiration Belt MR - Brain Products) and 2 EKG electrodes placed on the left subclavicular area and below the left axillary area. Of the twenty-five participants four were removed because of missing respiratory and/or cardiac data. The four audio stimuli come from https://www.franceinter.fr/emissions/une-histoire-et-oli : (1) Nadine et Robert les poissons rouges-Delphine de Vigan (8 min) (2) les villages du versant – Alice Zeniter (8 min) (3) Opaque et Opaline - Alex Vizorek (11 min) (4) le renard et le poulailler – Guillaume Meurice (10 min).

To test whether the ISC-HR is modulated by attention we divided the subjects into 2 groups: Group 1 (9 subjects) was recorded with stories (1) and (2) in a distracted condition, and (3) and (4) in an attentional condition. In group 2 (12 subjects) the stories were counterbalanced, stories (1) and (2) in the attentional condition, and (3) and (4) in the distracted condition. In the attentional condition (A), the subject’s task was to pay attention to the story while disregarding tones (320/360/400/440/482 Hz, 400ms long) that were played in random intervals (between 800ms and 1100ms) in the background of the story. After each story the subjects received a control debriefing questionnaire including 5 questions testing the memory performance of the story content. In the distracted condition (D), the subject’s task was to count backwards starting from 100 indexing the occurrence of ‘counting’ tones (same tones as in the attentional condition) in-between 2 ‘reset’ tones (Audacity- the type of tones is a linear decay between 1300 and 400 Hz during 400ms). The reset tones were added uniformly and randomly every 14 seconds on average. After each ‘reset’ tone, subjects had to reset the counting back to 100. At the end of the block the subjects had to report the smallest number obtained between 2 ‘reset’ tones. Subjects were instructed not to pay attention to the story and also receive the same debriefing questionnaire. The present research was promoted by the Inserm (CPP C13-41) and approved by the Comité de Protection des Personnes Ile-de-France 6. All subjects provided written informed consent.

#### Experiment 4: Auditory narratives with disorder of consciousness patients and healthy controls

Nineteen patients (8 females, age range 18 to 77, median age 50 years old) with disorders of consciousness (mostly resulting from brain lesions) and 24 healthy control subjects (14 females; age range 23-27, median 25 years old) listened to an auditory narrative (La part des ancêtres - Leonora Miano - 10 minutes, from: https://www.franceinter.fr/emissions/une-histoire-et-oli) through headphones while their EKG was recorded with a Polygraph Input Box (PIB of EGI-Geodesic’s physiological measurement system). The only instruction given to all subjects (healthy controls and patients) was to listen to the story. These patients were hospitalized in neurointensive care at Pitié Salpetrière (medical center with expertise in disorder of consciousness) to determine their state of consciousness, to adapt treatment, and to evaluate their neurological prognosis. During this evaluation, we performed several exams: clinical assessment, MRI, EEG, evoked response potential, and positron emission tomography. The state of consciousness is determined with some^61^ behavioral assessments, using the Coma Recovery Scale-revised^43^ - a score which allows differentiating between consciousness states: Coma (the patient does not open their eyes), Vegetative State (VS - Eye-opening, and alternance between wakefulness and sleep), Minimally Conscious State (MCS - the patient is able to follow their own face in the mirror or to follow a simple instruction) and Exit Minimally Conscious State (EMCS - patient can communicate with code). Among the 19 patients, we diagnosed 2 patients in coma, 8 VS patients, 8 MCS patients (7 MCS- and 1 MCS+) and 1 EMCS (see supplementary data for more details). The Ethical Committee of the Pitie-Salpetriere approved this research under the French label of ‘routine care research’.

### Computation of intersubject correlation of heart rate (ISC-HR)

Previous studies have relied on the quantification of synchrony of neuroimaging based time series (i.e. BOLD in fMRI ^32^ or signals from EEG electrodes ^26^). We follow a similar logic for the electrocardiographic (EKG) signal. We focus on the modulation of heart rate, by doing so we can determine if subjects increase or decrease their heart rate simultaneously, independently of their absolute level of heart rate. Step 1: We measure the EKG signal and detrend it using a high-pass filter (0.5 Hz cutoff) and subsequently notch filtered at either 50Hz (Experiment 1, 3 and 4) or 60 Hz (Experiment 2). We compute the instantaneous HR by detecting RR intervals from the EKG (Figure 1A). Peaks of the R-wave were found using *findpeaks* (built-in matlab function). Step 2: The instantaneous HR signal is then interpolated to keep the same sampling frequency for all subjects (Figure 1B). Step 3: This interpolated instantaneous HR signal is used to compute an inter-subject correlation matrix by calculating the Pearson’s correlation between each subjects’ instantaneous HR signal (Figure 1C). Step 4: Finally, the intersubject subject correlation of heart rate (ISC-HR) for each subject is obtained by computing the mean of correlations of that given subject to the rest of the group (Figure 1D).

For Experiment 2 (Fig 3. and 4) in step 3 we computed the inter-subject correlation matrix between the instantaneous HR signals in the attentive and distracted conditions with the instantaneous HR in the attentive conditions rather than within condition. In Experiment 3 (Fig. 5) we again used the instantaneous HR signals obtained when the groups were attentively listening to the stories as reference when computing the inter-subject correlation matrix for step 3 (Fig. 6.). For Experiment 4 we used the healthy participants as reference in the computation of the inter-subject correlation matrix in step 3. All other steps were as explained above.

### Statistical significance of ISC-HR

The instantaneous HR signals for a given epoch are first aligned across subjects. Then ISC-HR was calculated for all subjects in all epochs and the ISC across epochs. To test whether the ISC-HR value for each epoch was statistically significant (for Fig. 2A), circular shuffle statistic was used: Each subject’s instantaneous HR is circularly shifted by a random amount within the 60 second segments and the ISC-HR is re-computed. This procedure is repeated 10.000 times and the ISC-HR of the epoch is compared to this distribution of ISC-HR values for the circular shifted instantaneous HR signals. The p-value is obtained by counting how many circular shifted ISC-HR values were below the actual ISC-HR value. For Fig. 2B we repeat the circular shift to compute ISC-HR values and then average across epochs; p-values are then computed on these averaged ISC-HR values. For Figure 2C, instead of circular shifts within 60 s story segments, we instead randomly swapped segments between participants.

### Cluster statistics

One dimensional cluster statistics was computed by adapting the procedure described in ^79^ as follows. First, for each subject, we subtracted the spectrum in the attentive condition versus the distracted condition. Second, for each frequency we computed a t-test comparing the contrast to zero. Third, we identified clusters of consecutive frequencies with p-values < 0.05 and stored the sum of t-values within the cluster. Fourth, we run 10000 permutations in which we randomly reversed the sign of the subjects attentive versus distracted contrast and repeated steps 2-3 while keeping the sum of t-values of the largest cluster. Finally, we compared the clusters’ t-values obtained in step 3 with the distribution of permuted cluster t-values obtained in step 4. Clusters with larger than 95 % (corresponding to p-value<0.05) of the pemuted distribution were considered significant after multiple comparison cluster correction.

### Bayesian Factors

Bayes Factors are an established approach^80^ to compare the likelihood of a Null hypothesis to the likelihood of an alternative hypothesis expressing a measurable effect size. In our case, we were not able to measure a significant effect of synchronized breathing, and the question becomes whether there is sufficient evidence for a lack of synchronization, i.e. is there sufficient evidence in favour of the Null hypothesis. A Bayes Factor (BF) is the likelihood ratio between the Null and alternative hypothesis. When a BF lies between 3 and 10 it is considered moderate evidence in favour of the Null hypothesis. A crucial assumption when computing a BF is the effect size of the alternative hypothesis. If no specific effect size can be assumed a-priori, it is common to assume a distribution over a range of effect sizes. Rouder et al. formalizes this approach for the t-test.^81^ The conventional Null hypothesis for the t-test states that there is no difference between mean values of two groups and that mean values are normally distributed. For the alternative hypothesis Rouder assumes the Jeffreys-Zellner-Siow prior, which states that the effect size follows a Cauchy distribution, where effect size is the difference of means over the standard deviation. A similar default prior can be used for ANOVA^82^ and regression problems^83^. These default prior methods have the advantage that one can compute the BF directly from sample sizes and the t-statistics, F-statistics, or R-square, respectively. To compute a BF we use the matlab code of Bart Krekelberg, which implements these default priors (https://github.com/klabhub/bayesFactor).

### Frequency analysis of heart rate fluctuations

To investigate which time scale the inter-subject correlation and HRV is modulated by attention we do a frequency analysis of the instantaneous HR signal (Figure 4). The instantaneous HR signal was band-pass filtered using a 5th order butterworth filters with logarithmic spaced center frequencies with a bandwidth of 0.2 of the center frequency. The ISC was computed in each frequency band referenced to the attentive group (Fig. 4A). The HRV was computed as the standard deviation of the instantaneous HR normalized with the average HR (Fig. 4B).

### Computation of Fractional Anisotropy index (FA index)

Diffusion Tensor Images (DTI) were acquired on a 3T Siemens Skyra scanner (64 diffusion gradient directions, b value = 1000 s/mm^2^, TR/TE = 3000/80 ms, voxel size = 2×2×2 mm^3^). DTI data were pre-processed using the FDT package from the Functional MRI of the Brain (FMRIB) software library (FSL) package 5.01^84^. This consisted of: 1) correcting for motion and distortions caused by eddy currents; 2) brain segmentation using the brain extraction tool algorithm; 3) computing the fractional anisotropy (FA) maps using the diffusion-tensor model; 4) registration of the FA and MD maps on the FA template in the standard Montreal Neurological Institute (MNI) space using linear as well as nonlinear spatial transformations. FA values were averaged within a deep white-matter mask defined in the MNI space as the outline of the ICBM-DTI-81 white-matter labels atlas^85^. For each subject, this FA value was normalised with the mean of FA values measured from 10 healthy volunteers acquired with the same imaging protocol, such that an average FA index of 1.0 can be considered normal.

### Data availability

All healthy participants data and code to generate analysis and figures in this paper is accessible at https://osf.io/mhvy7/. In the case of patients, raw physiological data cannot be open due to consent limitations. In that case group data and code to generate the figures is available.

## Acknowledgements

This work was supported by Paris Brain Institute (France); Sorbonne PhD grant *(“recherches interdisciplinaires émergentes* “, France) to PP; CARNOT maturation grant, Title: Con&Heart to JDS; Sorbonne Universités EMERGENCE grant, title: Brain-body interactions, a new window for conscious evaluation in brain-injured patients to JDS; French embassy UK – Seeding Grant 2018 to JDS and DC; Medical Research Council New Investigator Research Grant (UK) to DC; National Science Foundation grant DRL-1660548 supported JM and LP.

## Supplementary materials

### Analysis of mean HR and HRV for experiment 1

For experiment 1 we also checked whether mean HR or HR variability (HRV) differed between story segments or between subjects (Fig. S1A and S1B). ANOVA with segment as fixed effect and subject as random effects shows that mean and standard deviation of HR differed between segments (HR: F(15,390)=2.46, p=1.89e-03, HRV: F(15,390)=1.73, p=4.26e-02), and between subjects (HR: F(26,390)=316.92, p=2.93e-244, HRV: F(26,390)=13.52, p=4.92e-40). We did not see a relationship between HR-ISC and mean HR (Fig. S1C, r(25)=0.31, p=0.11, BF01 = 1.95) or between HR-ISC and HRV (Fig. S1D, r(25)=-0.06, p=0.78, BF01 = 6.47).

**Supplementary Figure 1:**
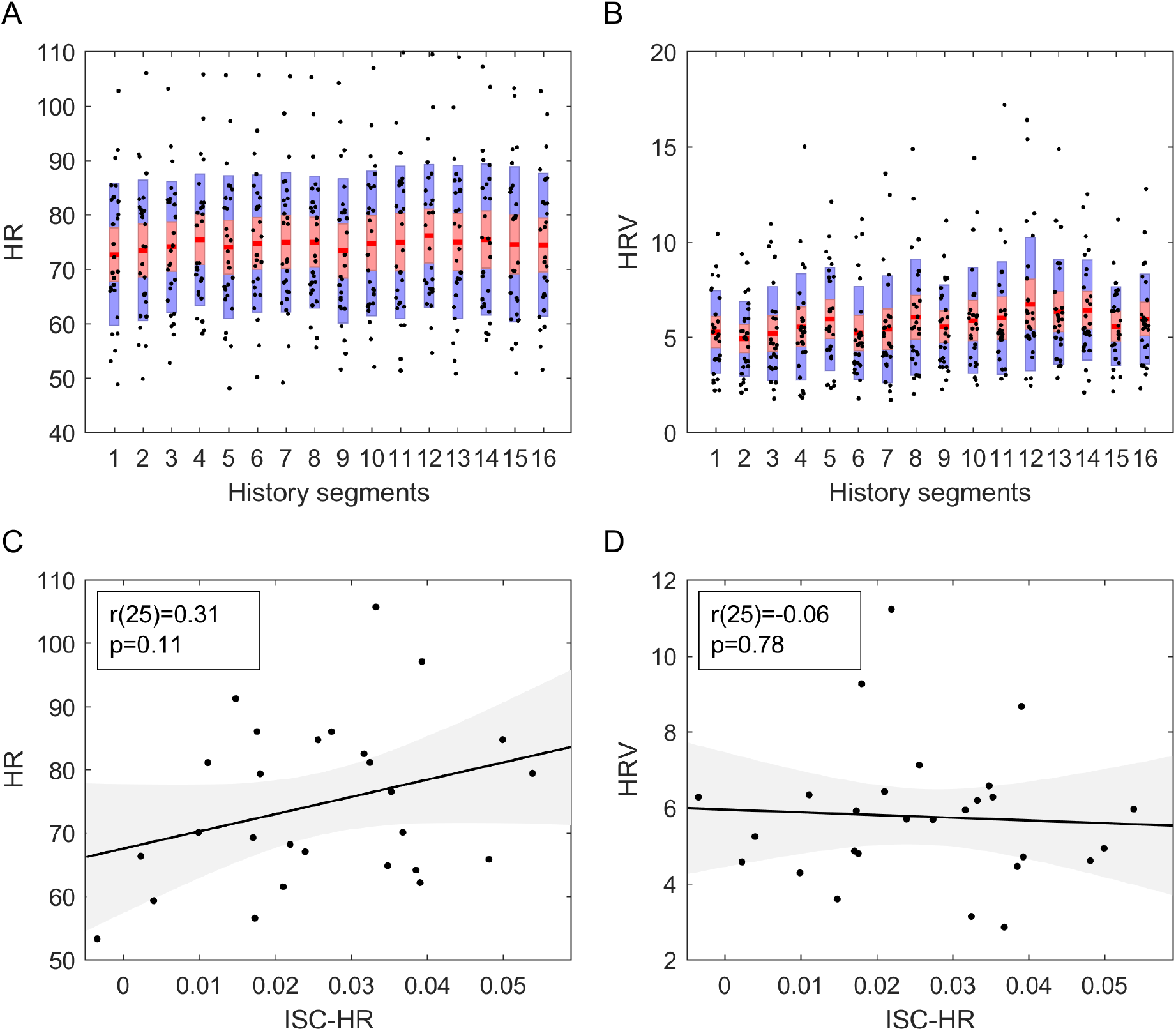
**A:** The instantaneous heart rate shows modulation across segments (Anova F(15,390) = 2.46, p=1.89e-03), and differences across subjects (Anova F(26,390)=316.92, p=2.93e-244). **B:** Heart rate variability measured as the standard deviation across the instantaneous heart rate for each subject (Anova F(26,390) = 13.52, p=4.92e-40), and differences across segments (Anova F(15,390)=1.73, p=4.26e-02). **C:** ISC-HR for each subject averaged across segments versus corresponding mean HR (each subject is a dot). We found no linear relationship between these two variables (BF01 = 1.95). **D:** ISC-HR for each subject averaged across segments versus corresponding HR variability (each subject is a dot). We found no linear relationship between these two variables (BF01 = 6.47).

### Analysis of mean HR and HRV for experiment 2

For experiment 2, in addition to the attentional effects reported in the main results section, we also checked whether mean HR or HR variability differed between each educational video and between subjects (Fig. S2A and S2B). ANOVA with video and conditions as fixed effect and subject as random effects shows that mean HR differed across videos (HR: F(4,211)=3.04, p=1.82e-02), but not standard deviation (F(4,211)=0.75, p=5.58e-01), and both differed significantly between subjects (HR: F(23,211)=89.26, p=1.74e-95, HRV: F(23,211)=29.61, p=1.39e-53). We did not see a relationship between ISC-HR and mean HR (Fig. S2C, Attentive: r(25)=-0.16, p=0.43, BF01 = 4.92, Distracted: r(25)=0.21, p=0.29, BF01 = 3.83) or between ISC-HR and HRV (Fig. S2D, Attentive: r(22)=0.15, p=0.48, BF01 = 4.96, Distracted: r(22)=0.46, p=0.02, BF10 = 1.95).

**Supplementary figure 2:**
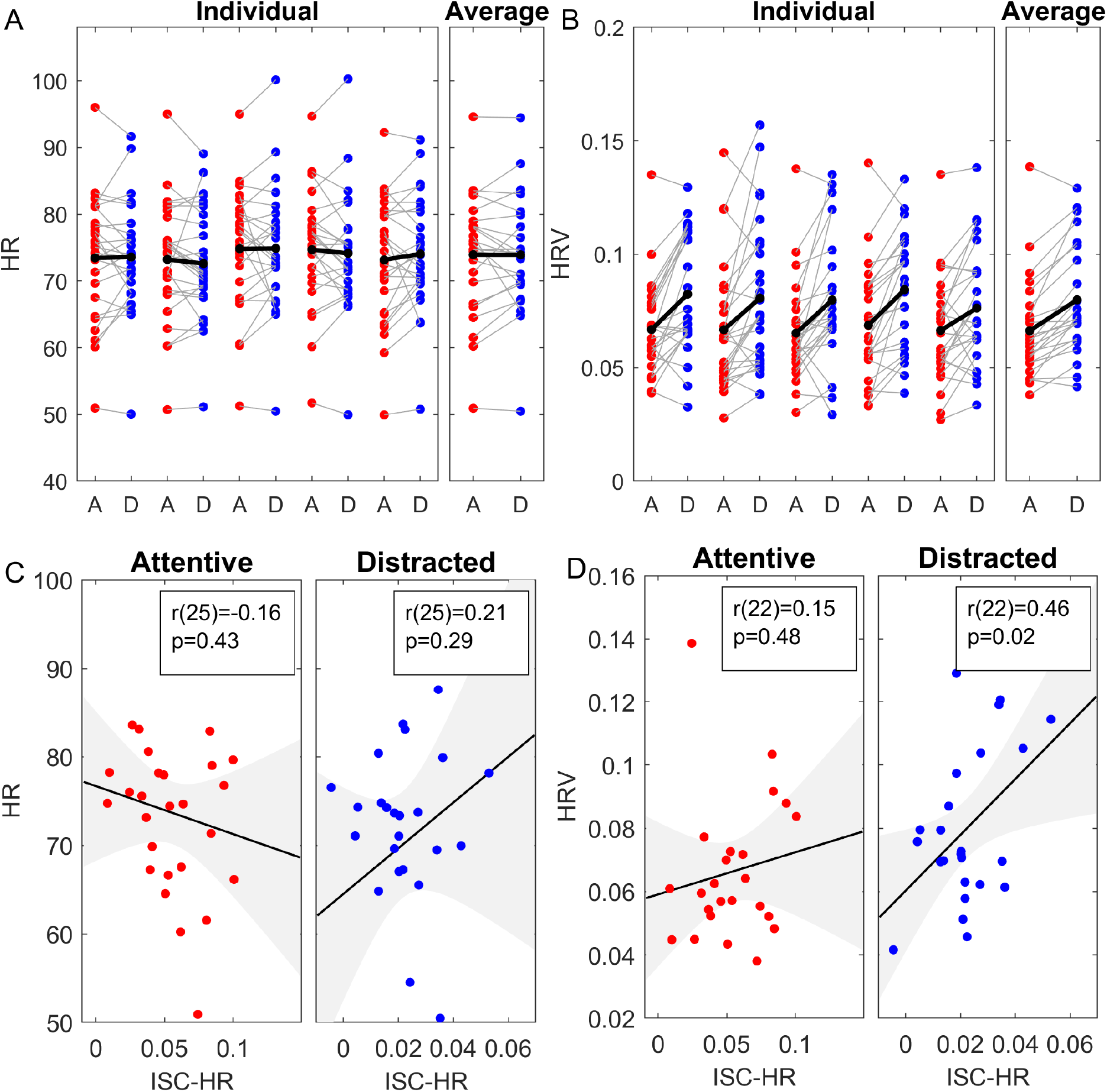
**A:** mean HR, **B:** HRV, **C**: correlation of ISC with HR. **D:** correlation of ISC with HRV

### Behavioral results of experiment 3

For experiment 3, we checked the modulation of memory performance in the attentive and distracted condition like a proof that the task was correctly completed. After each story, subjects answered 5 short questions about the story content testing their memory performance (i.e name of protagonist?, place?...). Memory performance was significantly higher in the attentive condition than in the distracted condition (Figure S3, Wilcoxon signed-rank test: z = 4.03, p = 5.7e-05, BF01 ~ 1e8). In none of the conditions we found a difference between group 1 and 2 in terms of memory performance. In both groups the performance was always similar, in the attentive condition (Median group 1 = 90; Median group 2 = 90; Mann-Whitney U test, U = 93, p = 0.67, BF01 = 2.3), in the distracted condition (Median group 1 = 40; Median group 2 = 30; Mann-Whitney U test, U = 106, p = 0.66, BF01 = 2.2), or in the contrast between the attentive and distracted conditions (Median group 1 = 40; Median group 2 = 65; Mann-Whitney U test, U = 86.5, p = 0.39, BF01 = 2.04)

**Supplementary figure 3:**
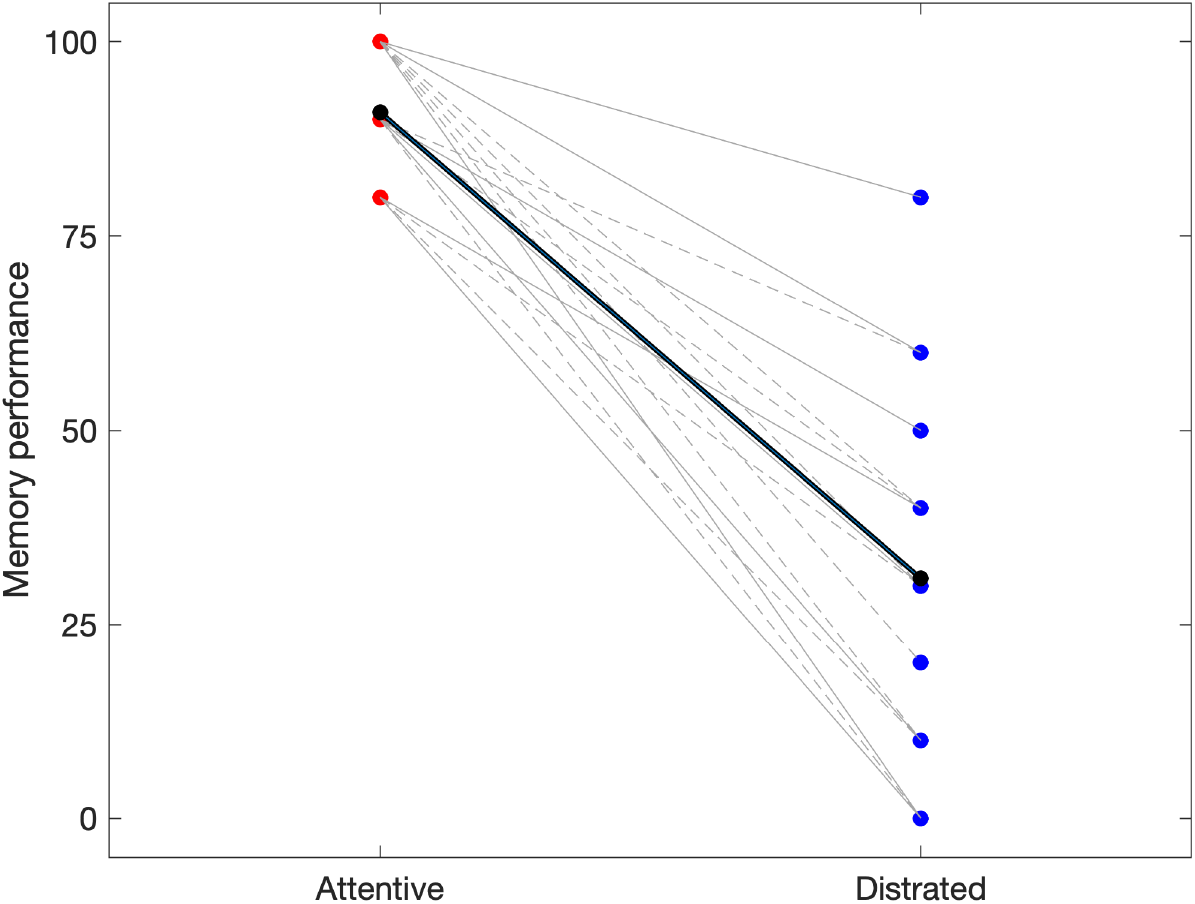
Behavioral responses attentive and distracted conditions (n=21). Solid lines correspond to subjects in group 1 and dashed lines to subjects in group 2. Each group listened to 2 two stories in the attentive condition and two in the distracted condition. Subjects answered 5 short questions about each story content. The memory performance (rate of good response) was systematically higher in the attentive condition compared to the distracted conditions. We observed no differences in memory performance between groups.

### Analysis of mean HR and HRV for experiment 3

For experiment 3 we also checked whether mean HR or HR variability differed between each condition and between subjects. We found that the mean HR was significantly higher in the distracted condition compared to the attentive condition (Fig S4A, paired t-test, t(20) = 7.32, p = 4.5e-7, BF10 = 37574). We found no differences in HRV between the attentive and distracted conditions (Fig S4B, paired t-test, t(20) = 1.64, p = 0.12, BF01 = 1.41).

We also did not see a relationship between ISC-HR in and mean HR (Fig. S4C, Attentive: r(19)=-0.08, p=0.74, BF01 = 5.67, Distracted: r(19)=0.22, p=0.33, BF01=3.74) or between ISC-HR and HRV (Fig. S4D, Attentive: r(19)=0.04, p=0.86, BF01 = 5.9, Distracted: r(19)=-0.14, p=0.55, BF01 = 5.01).

**Supplementary figure 4:**
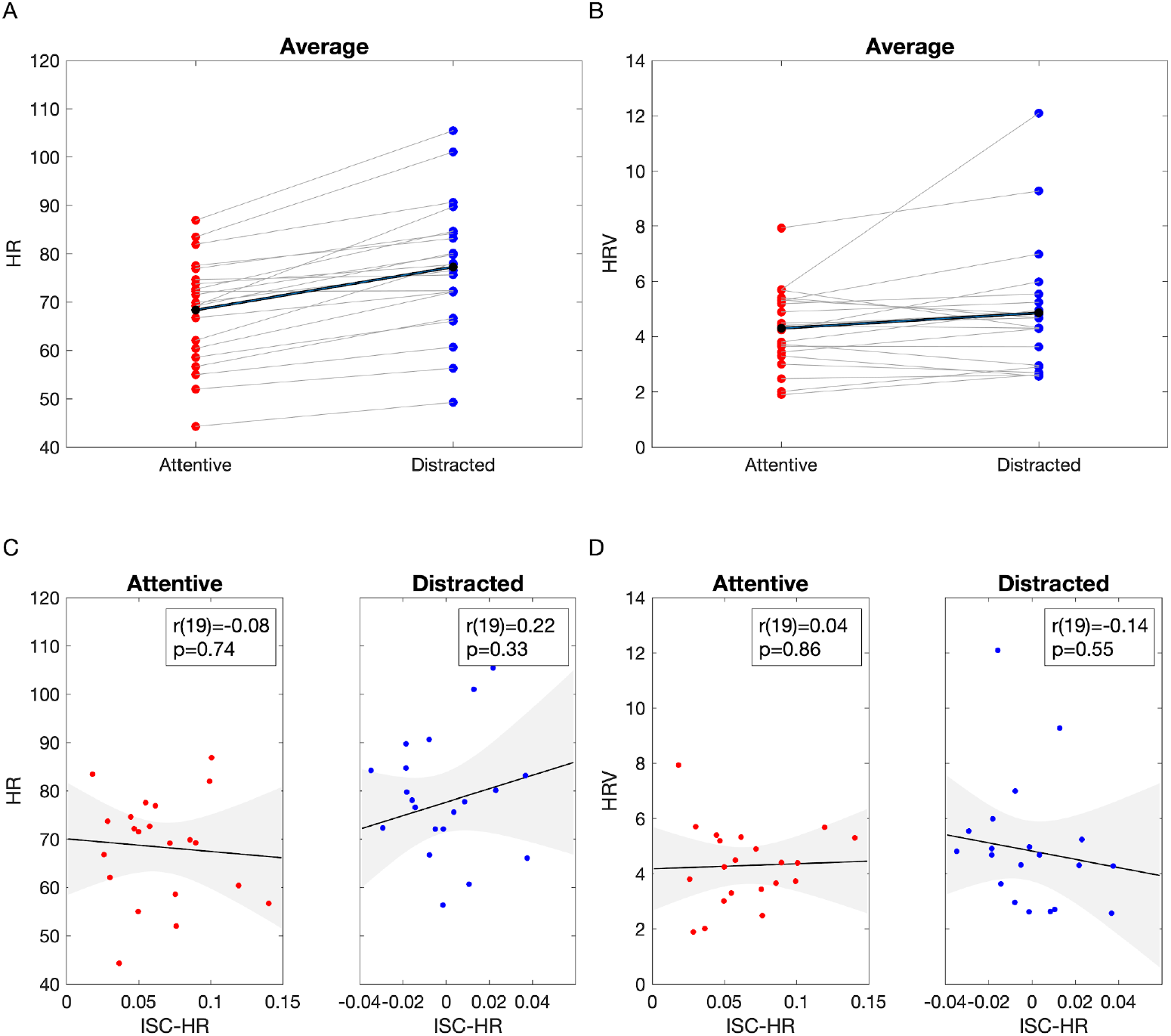
**A:** mean HR, **B:** HRV, **C**: correlation of ISC with HR in the attentive condition (left), and distracted condition (right), **D:** correlation of ISC with HRV in the attentive condition (left), and distracted condition (right)

### Respiration and HR and for experiment 3

Breathing is known to be a determinant of the heart rate. First, we tested whether attention affected the breathing signal itself. We computed the spectrum of the respiratory time-series in the attentive and distracted conditions (Fig. S5). We found two cluster corrected significant clusters, the first one from 0.34 Hz to 0.43 Hz (p = 0.0489) and from 0.63 Hz to 0.72 Hz (p = 0.0428).

Second, we extracted additional features: (1) cardio-respiratory coupling (Fig. S6A), (2) ISC computed from the raw respiratory signal (Fig S6B), (3) ISC computed from the instantaneous inspiratory frequency (Fig S6C), (4) ISC computed from the instantaneous expiratory frequency (Fig S6D), (5) ISC computed from the instantaneous inspiratory amplitude (Fig S6E), and (6) ISC computed from the instantaneous expiratory amplitude (Fig S6F). None of these respiratory features showed a significant ISC nor a modulation according to the attentional state (FDR corrected paired t-tests comparing the attentive and distracted conditions).

Statistical significance at the single-subject level of the computed features was determined by circular shuffling statistics.

**Figure S5:**
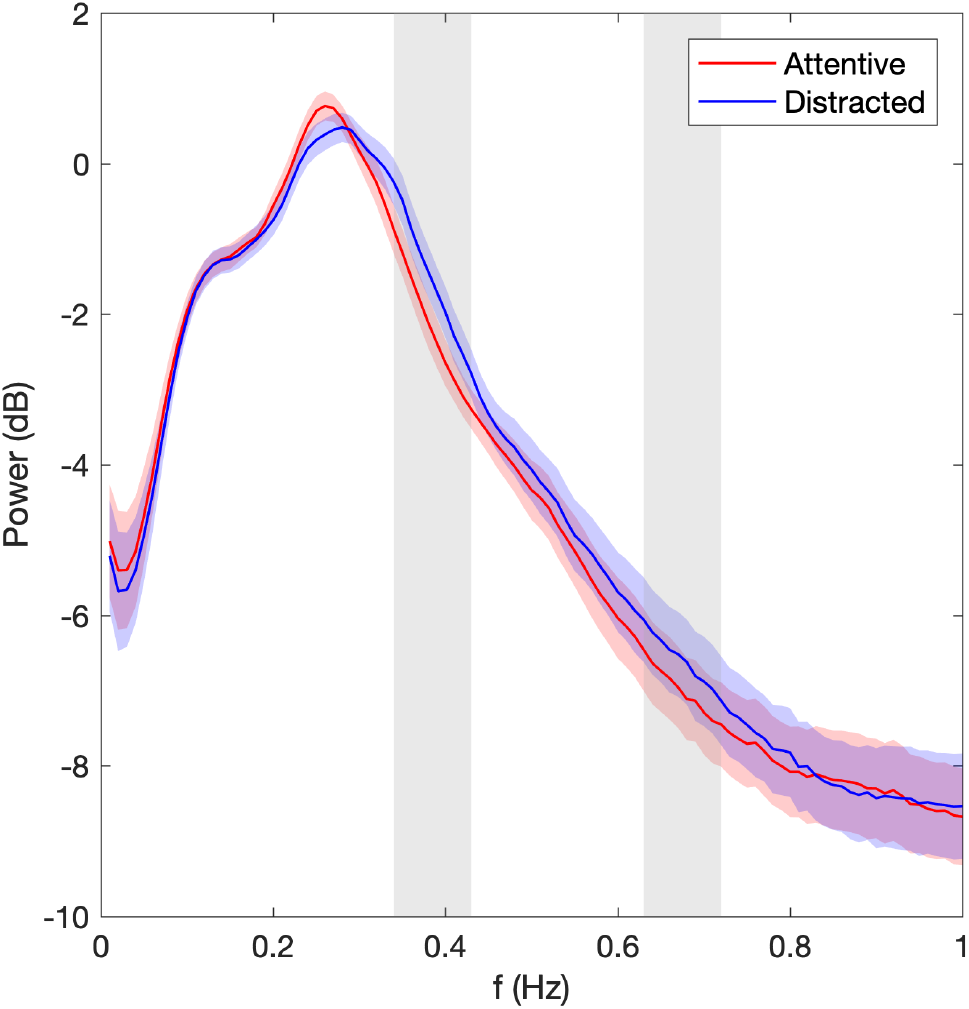
Power spectrum of breathing signal. multiple comparison corrected with onedimensional cluster statistics, p<0.05). Colored-shaded areas indicate standard error of the mean.

**Supplementary figure 6:**
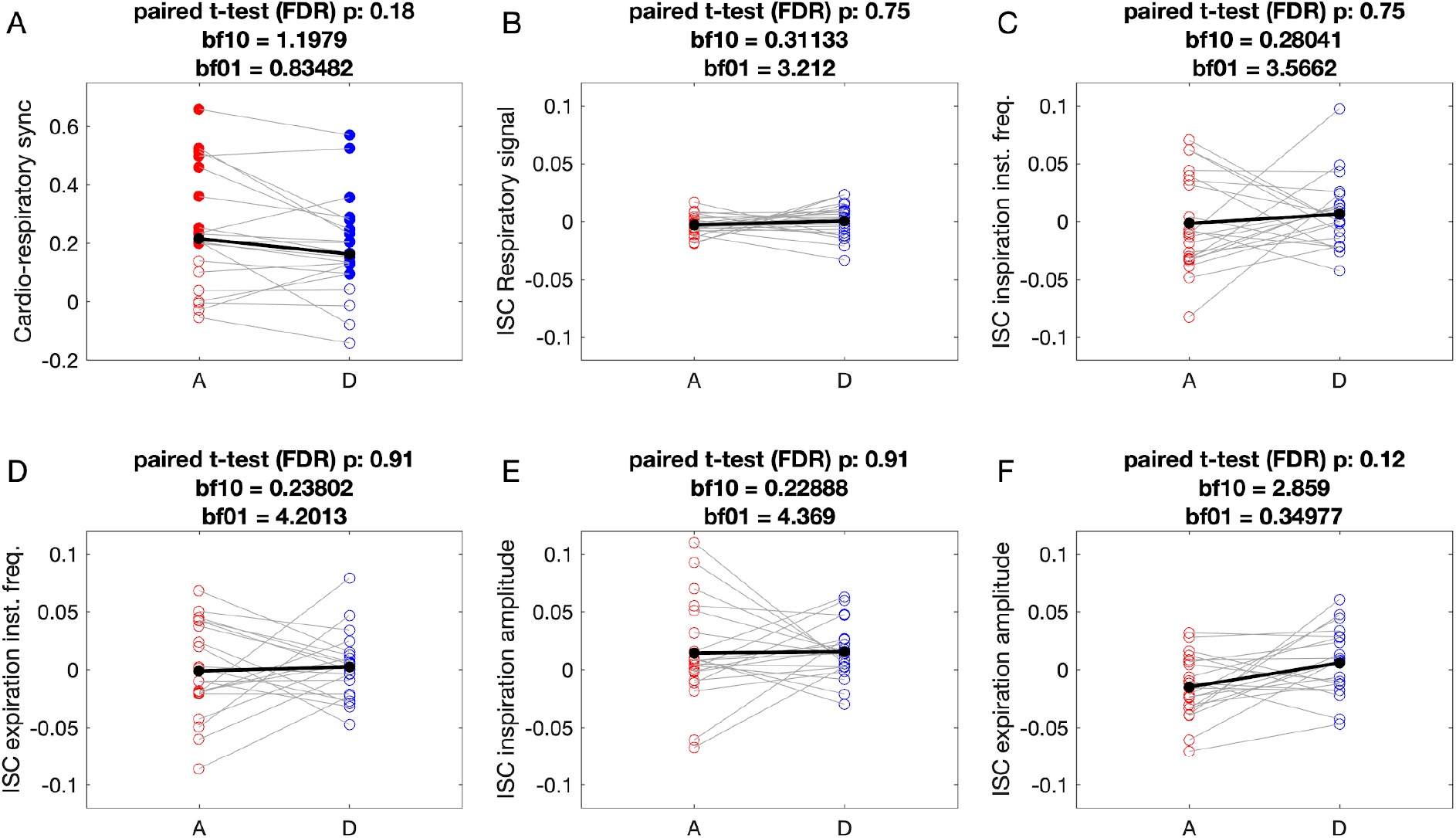
Attention modulation does not affect respiratory features synchronization. (A) Cardio-respiratory synchronization was significant in 12/21 subjects in the attentive condition and 15/21 in the distracted condition. However, we found no modulation of this synchrony with the attentional condition. (B-F) No individual significant ISC and no modulation of ISC with attention was found for raw respiratory signal (B), inspiratory instantaneous frequency (C), expiratory instantaneous frequency (D), inspiration amplitude (E), and expiration amplitude (F). Filled and empty circles indicate subject-level significant and nonsignificant respectively (FDR p<0.01 permutation statistics, circular shuffling statistics)

### Removal of respiration from heart rate signal

To remove any instantaneous or delayed effect of respiration on HR we estimate a linear impulse response from respiration to HR signals and subtract the HR signal estimated from the respiration using this impulse response. The residual HR signal is guaranteed to be uncorrelated from respiration at any delay. The filter is estimated separately for each subject and attention condition using 20s of data and ISC is computed on the remaining data using the residual HR signal. The ISC results reported in the main paper (Fig. 5) are reproduced here for the residual HR signal (Supplementary Fig. 6). With the exception of one outlier subject in the distracted conditions, the modulation with attention is essentially unchanged (t-test: t(20) = 5.57, p = 1.9e-05, signrank: p = 0.00048, BF = 1251.3)

**Supplementary figure 7:**
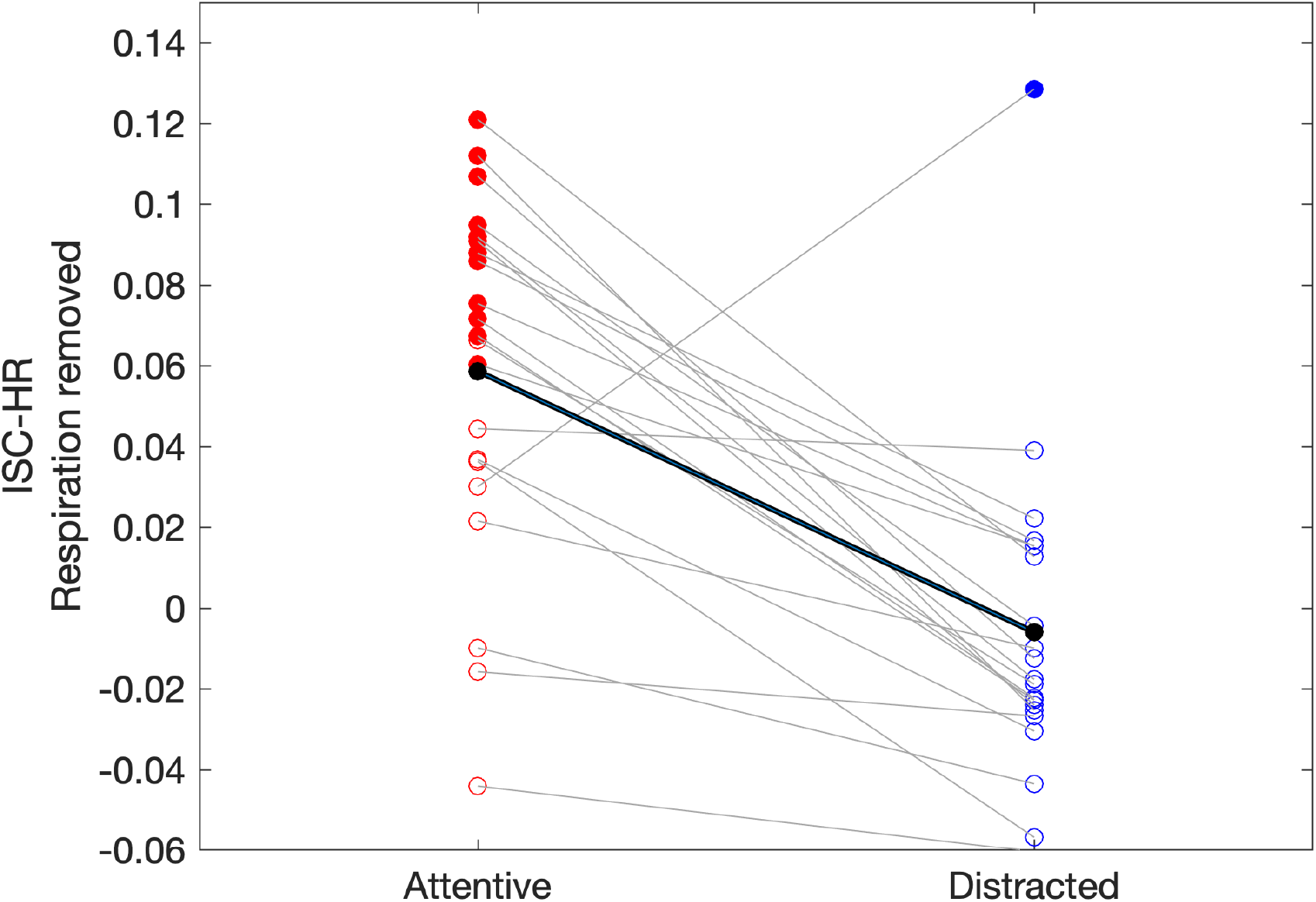
Same as Fig. 5A but computed after removing the effect of respiration on HR allowing for potentially delayed correlation. Filled and empty circles indicate subject-level significant and non-significant respectively (FDR p<0.01 permutation statistics, circular shuffling statistics)

### Respiration and HR and for experiment 4

In experiment 4 we replicated the analysis comparing cardiac and respiratory features in healthy participants. We computed the ISC from the: raw respiration (RR), instantaneous inspiratory rate (IIR), instantaneous expiratory rate (IER), inspiratory amplitude (IA), expiratory amplitude (EA), instantaneous heart rate (IHR), (7) instantaneous heart rate after respiratory removal (IHR_r). While none of the respiratory ISCs show a significant difference to zero at the group level, both HR-ISC were different to zero (Table 1).

In addition, both IHR and IHR_r were systematically higher than the ISC computed from respiratory features (Table 2).

**Table 1.** Experiment 4, respiratory and cardiac features stats at the group level compared to zero.

| ISC |  | T-stats | p-value | p-value (FDR) | BF10 | BF01 |
| --- | --- | --- | --- | --- | --- | --- |
| Resp. | Raw Signal | $t(22) = 0.64$ | 0.53 | 0.53 | 0.26 | 3.86 |
| | Inst. Insp. Rate | $t(22) = 1.80$ | 0.08 | 0.17 | 0.86 | 1.16 |
| | Inst. Exp. Rate | $t(22) = 1.72$ | 0.10 | 0.17 | 0.77 | 1.30 |
| | Insp. Amplitude | $t(22) = 1.27$ | 0.22 | 0.25 | 0.44 | 2.27 |
| | Exp. Amplitude | $t(22) = -1.51$ | 0.14 | 0.20 | 0.58 | 1.71 |
| Heart | Inst. HR | $t(23) = 6.06$ | $3.52e-6$ | $2.01e-5$ | 5636 | $1.7e-4$ |
| | Inst. HR (-resp) | $t(22) = 5.93$ | $5.75e-6$ | $2.01e-5$ | 3635 | $2.7e-4$ |

**Table 2.** Experiment 4, comparison between respiratory and cardiac features.

|  | Instant. HR |  |  |  |  | Instant. HR (-resp) |  |  |  |  |
| --- | --- | --- | --- | --- | --- | --- | --- | --- | --- | --- |
|  | T-stats | p-value | p-val (FDR) | BF10 | BF01 | T-stats | p-value | p-val (FDR) | BF10 | BF01 |
| Raw Signal | $t(22) = 5.24$ | $2.9e-5$ | $7.3e-5$ | 823 | 0.001 | $t(22) = 5.33$ | $2.37e-5$ | $5.93e-5$ | 1001 | 0.001 |
| Inst. Insp. Rate | $t(22) = 2.20$ | 0.04 | 0.04 | 1.61 | 0.62 | $t(22) = 2.83$ | 0.01 | 0.01 | 5.0 | 0.2 |
| Inst. Exp. Rate | $t(22) = 2.81$ | 0.01 | 0.01 | 4.83 | 0.20 | $t(22) = 3.5$ | 0.002 | 0.003 | 19 | 0.05 |
| Insp. Amplitude | $t(22) = 2.78$ | 0.01 | 0.01 | 4.57 | 0.21 | $t(22) = 3.20$ | 0.004 | 0.005 | 10.3 | 0.1 |
| Exp. Amplitude | $t(22) = 5.35$ | $2.27e-5$ | $7.36e-5$ | 1044 | 0.001 | $t(22) = 5.6$ | $1.2e-5$ | $5.9e-5$ | 1807 | 0.0005 |

**Table 3.** Description of patient cohort: summary table including patients with sex, age, etiology of brain lesion, delay between brain lesion and the evaluation, state of consciousness, CRS-R (Coma Recovery Scale-Revised), Fractional anisotropy (FA) from MRI and outcome. *Outcome collected before 6 months.

| Subject | Sex | Age (years) | Etiology | Delay (days) | State | CRS-R | FA | Outcome |
| --- | --- | --- | --- | --- | --- | --- | --- | --- |
| 1 | M | 71 | Acute Polyradiculoneuritis | 12 | Coma | 0 | no exam | VS/Death* |
| 2 | F | 77 | Meningitis | 86 | Coma | 2 | no exam | VS |
| 3 | M | 18 | TBI+Anoxia | 170 | VS | 5 | 0.756 | VS |
| 4 | M | 29 | TBI+Anoxia | 510 | VS | 5 | artefacted | VS |
| 5 | M | 29 | Anoxia | 556 | VS | 5 | 0.492 | VS |
| 6 | M | 55 | TBI | 27 | VS | 6 | artefacted | MCS+ |
| 7 | F | 53 | Anoxia | 328 | VS | 5 | 0.824 | VS |
| 8 | F | 24 | TBI | 68 | VS | 5 | 0.644 | MCS- |
| 9 | F | 63 | Anoxia | 41 | VS | 5 | 0.642 | VS |
| 10 | M | 18 | Anoxia | 59 | VS | 4 | 0.792 | VS/Death* |
| 11 | F | 50 | Anoxia | 5852 | MCS- | 10 | 0.698 | MCS- |
| 12 | M | 26 | Status Epilepticus | 209 | MCS- | 9 | 0.561 | MCS+ |
| 13 | M | 50 | Anoxia | 60 | MCS- | 7 | 0.85 | EMCS |
| 14 | F | 61 | Status Epilepticus | 90 | MCS- | 10 | noDTI | MCS- |
| 15 | M | 58 | Anoxia | 302 | MCS- | 14 | 0.548 | MCS- |
| 16 | M | 56 | TBI | 478 | MCS- | 11 | 0.824 | MCS-* |
| 17 | F | 36 | Left hematoma | 121 | MCS- | 12 | noDTI | EMCS |
| 18 | F | 49 | TBI | 980 | MCS+ | 13 | noDTI | MCS+ |
| 19 | M | 28 | Status Epilepticus | 313 | EMCS | 14 | no exam | EMCS |

Finally, IHR_r was significantly higher than IHR (p-value (t-test, FDR) = 0.006, BF10 = 7.3)

**Supplementary figure 8:**
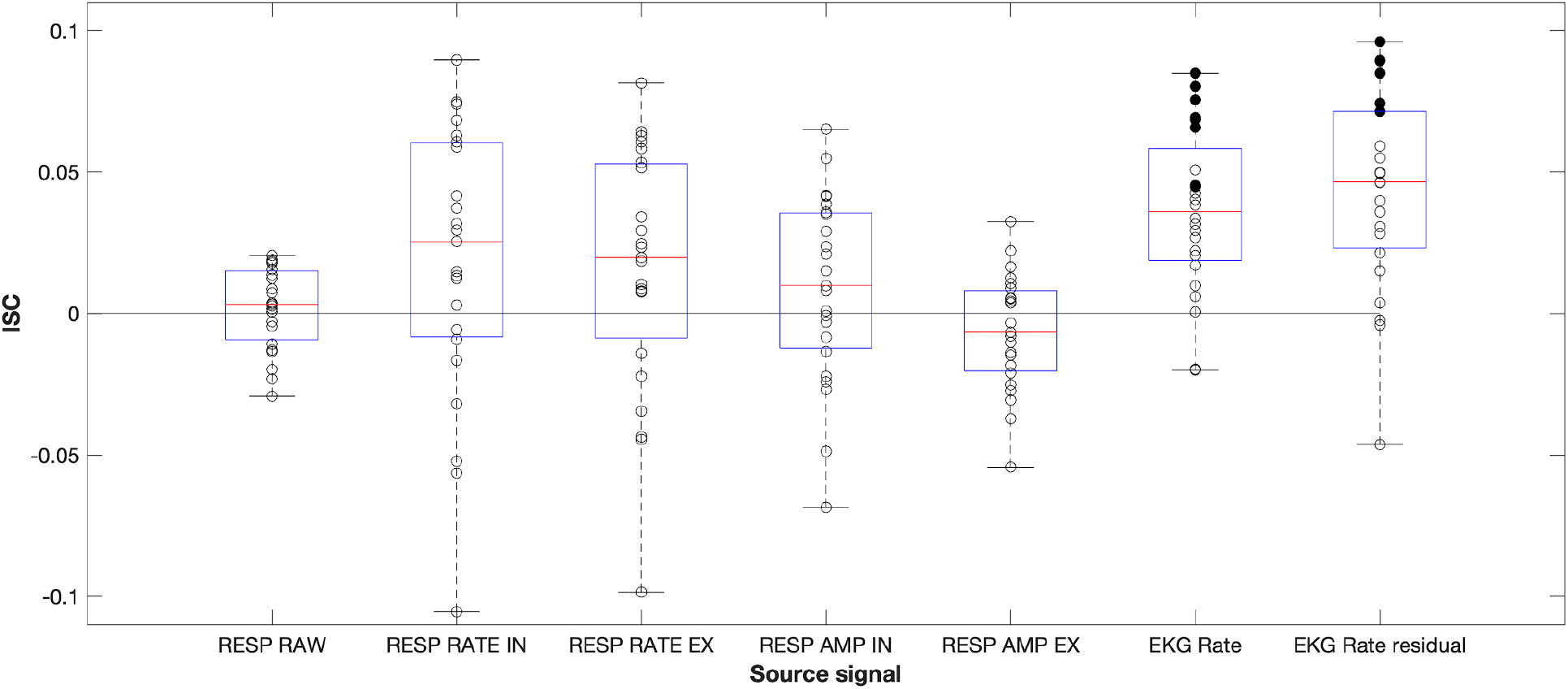
EKG-ISC synchronization outperformed respiratory features synchronization. Filled and empty circles indicate subject-level significant and non-significant respectively (FDR p<0.01 permutation statistics, circular shuffling statistics)

### Analysis of mean HR and HRV for experiment 4

For experiment 4 we also checked whether mean HR or HR variability differed between the different states of consciousness (Fig. S9). The mean HR of DoC patients was significantly higher than healthy participants (Fig S9A, t(41) = 4.7, p = 2.9e-05, BF10 = 12.4). To test if HR changed with the state-of-consciousness among the DoC patients we performed a Spearman correlation between state-of-consciousness (Coma = 1, VS = 2, MCS- = 2, MCS+ = 3, EMCS = 4) and HR and found no significant correlation (R(17) = −0.18, p = 0.45, BF01 = 3.37). Similar for the HR variability, HRV was significantly higher in DoC patients compared to healthy controls (Fig S6B, t(41) = 2.34, p = 0.02, BF10 = 2.54). We found no significant relationship between the patients’ state-of-consciousness and HRV (Spearman correlation, R(17) = 0.41, p = 0.08, BF01 = 2.11).

We also controlled the relationships between HR and ISC, and HRV and ISC for patients. In none of the cases the relationship was significant (Fig. S9C, HR versus ISC, r(17)=0.08, p=0.74; BF01 = 5.42, Fig. S9D, r(17)=-0.4, p=0.09, BF01 = 1.45).

**Supplementary figure 9:**
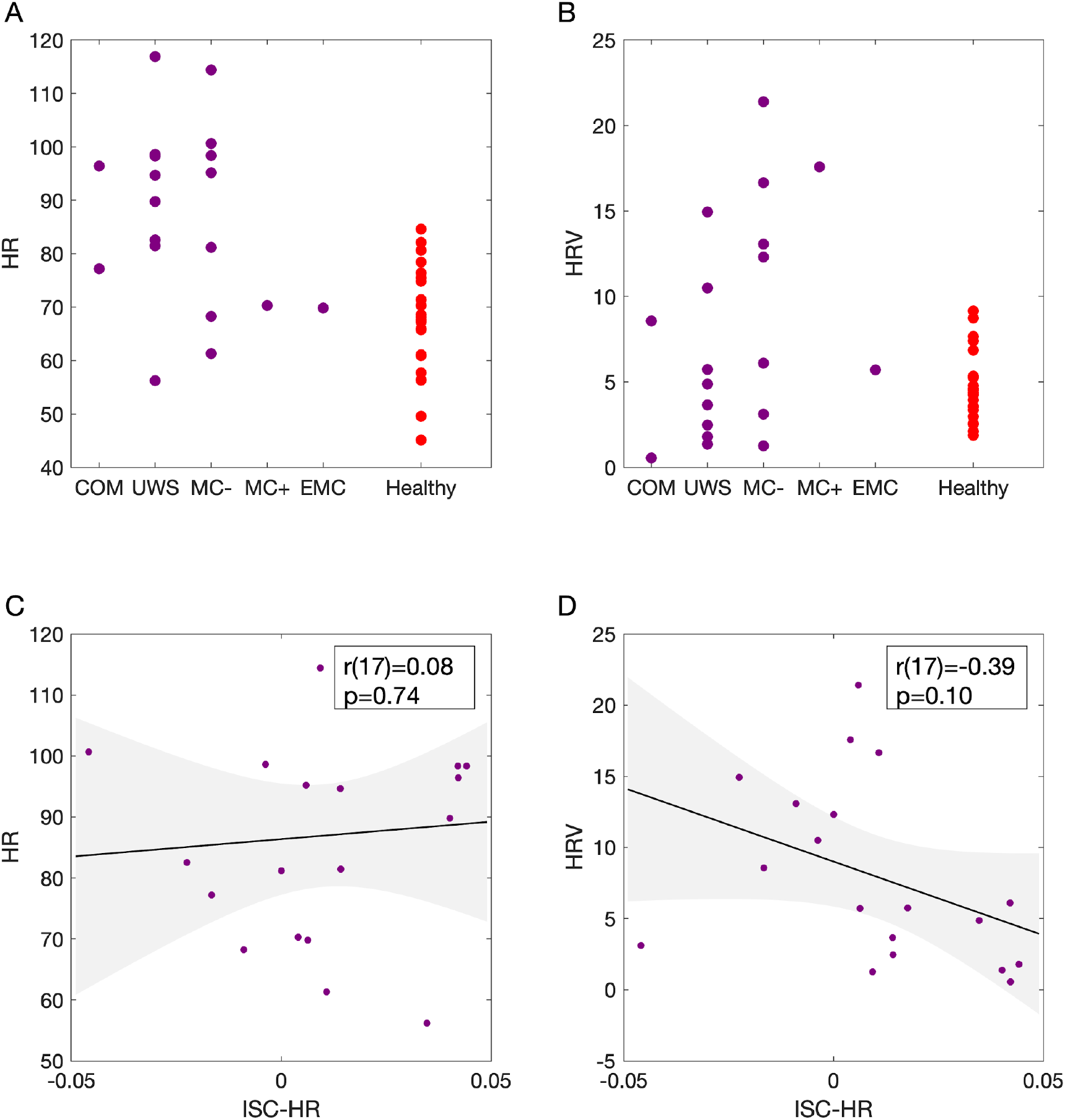
**A:** mean HR for DoC patients and healthy participants (Note that DoC patients are ordered in individual columns corresponding to their clinical state-of-consciousness (1=Coma, 2=VS/UWS, 3=MCS-, 4=MCS+, and 5=EMCS) **B:** HRV, idem to A **C**: correlation of ISC with HR for DoC patients. **D:** correlation of ISC with HRV for DOC patients.

